# Changes in Forest Composition, Stem Density, and Biomass from the Settlement Era (1800s) to Present in the Upper Midwestern United States

**DOI:** 10.1101/026575

**Authors:** Simon J. Goring, David J. Mladenoff, Charles V. Cogbill, Sydne Record, Christopher J. Paciorek, Stephen T. Jackson, Michael C. Dietze, Andria Dawson, Jaclyn Hatala Matthes, Jason S. McLachlan, John W. Williams

**Author notes:** **Corresponding Author**: Simon Goring.

## Abstract

EuroAmerican land use and its legacies have transformed forest structure and composition across the United States (US). More accurate reconstructions of historical states are critical to understanding the processes governing past, current, and future forest dynamics. Gridded (8×8km) estimates of pre-settlement (1800s) forests from the upper Midwestern US (Minnesota, Wisconsin, and most of Michigan) using 19th Century Public Land Survey (PLS) records provide relative composition, biomass, stem density, and basal area for 26 tree genera. This mapping is more robust than past efforts, using spatially varying correction factors to accommodate sampling design, azimuthal censoring, and biases in tree selection.

We compare pre-settlement to modern forests using Forest Inventory and Analysis (FIA) data, with respect to structural changes and the prevalence of lost forests, pre-settlement forests with no current analogue, and novel forests, modern forests with no past analogs. Differences between PLSS and FIA forests are spatially structured as a result of differences in the underlying ecology and land use impacts in the Upper Midwestern United States. Modern biomass is higher than pre-settlement biomass in the northwest (Minnesota and northeastern Wisconsin, including regions that were historically open savanna), and lower in the east (eastern Wisconsin and Michigan), due to shifts in species composition and, presumably, average stand age. Modern forests are more homogeneous, and ecotonal gradients are more diffuse today than in the past. Novel forest assemblages represent 29% of all FIA cells, while 25% of pre-settlement forests no longer exist in a modern context.

Lost forests are centered around the forests of the Tension Zone, particularly in hemlock dominated forests of north-central Wisconsin, and in oak-elm-basswood forests along the forest-prairie boundary in south central Minnesota and eastern Wisconsin. Novel FIA forest assemblages are distributed evenly across the region, but novelty shows a strong relationship to spatial distance from remnant forests in the upper Midwest, with novelty predicted at between 20 to 60km from remnants, depending on historical forest type.

The spatial relationships between remnant and novel forests, shifts in ecotone structure and the loss of historic forest types point to significant challenges to land managers if landscape restoration is a priority in the region. The spatial signals of novelty and ecological change also point to potential challenges in using modern spatial distributions of species and communities and their relationship to underlying geophysical and climatic attributes in understanding potential responses to changing climate. The signal of human settlement on modern forests is broad, spatially varying and acts to homogenize modern forests relative to their historic counterparts, with significant implications for future management.

## Introduction

The composition, demography, and structure of forests in eastern North America have changed continuously over the last millennium, driven by human land use [1–5] and climate variability [6–9] While human effects have been a component of these systems for millenia, the EuroAmerican settlement and industrialization period have increased anthropogenic effects by orders of magnitude [10–12]. Legacies of post-settlement land use in the upper Midwest [13] and elsewhere have been shown to persist at local and regional scales [5,14,15], and nearly all North American forests have been affected by the intensification of land use in the past three centuries. Hence, contemporary ecological processes in North American forests integrate the anthropogenic impacts of the post-EuroAmerican period and natural influences at decadal to centennial scales.

At a regional scale many forests in the upper Midwest (*i.e*., Minnesota, Wisconsin and Michigan) now have decreased species richness and functional diversity relative to forests of the pre-EuroAmerican settlement (hereafter pre-settlement) period [16–18] due to near complete logging. For example, forests in Wisconsin are in a state of regrowth, with an unfilled carbon sequestration potential of 69 TgC [19] as a consequence of these extensive land cover conversions and subsequent partial recovery following abandonment of farm lands in the 1930s. But while regional patterns may establish themselves across the midwest, the range of ecozones and patterns of land use in space and time result in both broad spatial patterns, but significant local to regional variation. For example, while fire suppression occured throughout the region effects of suppression have and will continue to manifest themselves differently depending on the historical vegetation and biophyssical characteristics of the site or region.

Legacies of land use are unavoidable at regional scales [20]. Under intensive land use change the natural processes of secession, senescense and the replacement of tree species in forests may be masked, or heavily modified by historically recent land use change. Broad-scale land use change can result in changes to forest structure and species pools that may result in non-stationarity within ecosystems that may not be apparent on the relatively narrow time scales at which ecology traditionally operates [21], meaning chronosequences may not be sufficeint to understand shifts in structure and composition. There is a history of recolonization of forested landscapes following agricultural clearance in the upper Midwest [22], pointing to the importance of understanding ecological trajectories and land use legacies in understanding modern forest dynamics [20]. Cramer*el al*.. [23] point to the literature of succession theory to indicate the likelihood that many old fields will return to a ‘natural’ state, but point out that recovery is not universal. In particular, intense fragmentation of the landscape can deplete the regional species pool, leading to failures of recruitment that would favor species with longer distance seed dispersersal [24]. In the upper Midwest long seed dispersal would favor species such as poplar (*Populus* sp.), white birch (*Betula papyrifera*) and some maple species (*Acer* sp.), at the expense of large-seeded species such as walnut (*Juglans* sp.), oak (*Quercus* sp.) and others.

While there remains debate over the utility of the concept of novel ecosystems [25,26], the fact remains that there are now forest and vegetation communities on the landscape without past analogues. The long term management of the systems and their associated services requires a broad understanding of the extent to which landscapes have been modified, and the extent to which land use change has potenitally masked underlying processes. It also requires a better understanding of the spatial (and temporal) scales at which novel ecosystems operate. While restoration efforts have generally focused on ecosystems at local scales, there is an increasing need to focus on management and restoration at landscape scales [27]. Thus a better understanding of the landscape-scale processes driving novelty, the spatial structure of novel ecosystems and their ecological correlates, is increasingly important. An understanding of landscape level processes driving ecological novelty can help prioritize intervention strategies at local scales [28], and give us a better understanding of the role of patches in restoring hybrid or novel landscapes. In particular, how important is the species pool to the development of novel landscapes? Are novel forests further from remnant forests than might otherwise be expected? Is novelty operating at landscape scales in the upper Midwest, and is the spatial distribution of new forests tied to historical patterns vegetation or losses of forest types from the historical landscape?

The upper Midwestern United States represents a unique ecological setting, with multiple major ecotones, including the prairie-forest boundary, historic savanna, and the Tension Zone between southern deciduous forests and northern evergreen forests. The extent to which these ecotones have shifted, and their extent both prior to and following EuroAmerican settlement is of critical importance to biogeochemical and biogeophysical vegetation-atmosphere feedbacks [29], carbon sequestration [19], and regional management and conservation policy [30–33].

Land use change at the local and state-level has affected both the structure and composition of forests in the Midwestern United States [16,17]. Homogenization and shifts in overall forest composition are evident, but the spatial extent and structure of this effect is less well understood. Studies in Wisconsin have shown differential patterns of change in the mixedwood and evergreen dominated north, the southern driftless and hardwood dominated forests in south-central Wisconsin, and the prairie and savanna ecosystems that bound the region to the south and west. Does this pattern of differential change extend to Minnesota and Michigan? To what extent are land-use effects common across the region, and where are responses ecozone-specific? Has homogenization [16] resulted in novel forest assemblages relative to pre-settlement baselines across the region, and the loss of pre-settlement forest types? Are the spatial distributions of these novel and lost forest types overlapping, or do they have non-overlapping extents? If broad-scale reorganization is the norm following EuroAmerican settlement, then the ecosystems that we have been studying for the past century may indeed be novel relative to the reference conditions of the pre-settlement era.

Modern forest structure and composition data [34] play a ubiquitous role in forest management, conservation, carbon accounting, and basic research on forest ecosystems and community dynamics. These recent surveys (the earliest FIA surveys began in the 1930s) can be extended with longer-term historical data to understand how forest composition has changed since EuroAmerican settlement. The Public Land Survey was carried out ahead of mass EuroAmerican settlement west and south of Ohio to provide for delineation and sale of the public domain beyond the original East Coast states [35,36]. Because surveyors used trees to locate survey points, recording the identity, distance, and directory of two to four trees next to each survey marker, we can make broad-scale inferences about forest composition and structure in the United States prior to large-scale EuroAmerican settlement [37–40]. In general, FIA datasets are systematically organized and widely available to the forest ecology and modeling community, whereas most PLSS data compilations are of local or, at most, state-level extent. This absence of widely available data on settlement-era forest composition and structure limits our ability to understand and model the current and future processes governing forest dynamics at broader, regional scales. For example, distributional models of tree species often rely upon FIA or other contemporary observational data to build species-climate relationships that can be used to predict potential range shifts [41,42].

Here we use survey data from the original Public Lands Surveys (PLS) in the upper Midwest to derive estimates of pre-settlement (*ca*. mid-late 1800s) forest composition, basal area, stem density, and biomass. This work builds upon prior digitization and classification of PLSS data for Wisconsin [43,44] and for parts of Minnesota [17,45] and Michigan Michigan (USFS-NCRS http://www.ncrs.fs.fed.us/gla/). Most prior PLS-based reconstructions are for individual states or smaller extents [17,19,45,46] often aggregated at the scale of regional forest zones [16,17], although aggregation may also occur at the section [19] or township scale [47]. Our work develops new approaches to address major challenges to PLSS data, including lack of standardization in tree species names, azimuthal censoring by surveyors, variations in sampling design over time, and differential biases in tree selection among different kinds of survey points within the survey design at any point in time. The correction factors developed here are spatially varying, allowing us to accommodate temporal and spatial variations in surveyor methods.

We aggregate point based estimates of stem density, basal area and biomass to an 8 x 8km grid, and classify forest types in the upper Midwest to facilitate comparisons between FIA and PLSS data. We compare the PLSS data to late-20th-century estimates of forest composition, tree stem density, basal area and biomass. We explore how forest homogenization has changed the structure of ecotones along two major ecotones from southern deciduous to northern evergreen forests and to the forest-prairie boundary. Using analog analyses, we identify lost forests that have no close compositional counterpart today and novel forests with no close historical analogs. This work provides insight into the compositional and structural changes between historic and contemporary forests, while setting the methodological foundation for a new generation of maps and analyses of settlement-era forests in the Eastern US.

## Methods

### Public Lands Survey Data: Assembly, and Standardization

The PLSS was designed to facilitate the division and sale of federal lands from Ohio westward and south The survey created a 1 mile^2^ (2.56 km^2^) grid (sections) on the landscape. At each section corner, a stake was placed as the official location marker. To mark these survey points, PLSS surveyors recorded tree stem diameters, measured distances and azimuths of the two to four trees ‘closest’ to the survey point and identified tree taxa using common (and often regionally idiosyncratic) names. PLSS data thus represent measurements by hundreds of surveyors from 1832 until 1907, with changing sets of instructions over time (Stewart, 1979).

The PLSS was undertaken to survey land prior to assigning ownership (Stewart 1935, White 1983), replacing earlier town proprietor surveys (TPS) used for the northeastern states [2,48]. The TPS provided estimates of relative forest composition at the township level, but no structural attributes. The PLSS produced spatially explicit point level data, with information about tree spacing and diameter, that can be used to estimate absolute tree density and biomass. PLSS notes include tree identification at the plot level, disturbance [49] and other features of the pre-settlement landscape. However, uncertainties exist within the PLSS and township level dataset [50].

Ecological uncertainty in the PLSS arises from the dispersed spatial sampling design (fixed sampling every 1 mile), precision and accuracy in converting surveyor’s use of common names for tree species to scientific nomenclature [51], digitization of the original survey notes, and surveyor bias during sampling [38,50,52,53]. Estimates vary regarding the ecological significance of surveyor bias. Terrail *et al.* [54] show strong fidelity between taxon abundance in early land surveys versus old growth plot surveys. Liu *et al* [38] estimate the ecological significance of some of the underlying sources of bias in the PLSS and show ecologically significant (>10% difference between classes) bias in species and size selection for corner trees. However Liu *et al.* [38] also indicate that the true sampling error cannot be determined, particularly because most of these historic ecosystems are largely lost to us.

Kronenfeld and Wang [55], working with historical land cover datasets in western New York indicate that direct estimates of density using plotless estimators may be off by nearly 37% due to azimuthal censoring (i.e., the tendency of surveyors to avoid trees close to cardinal directions), while species composition estimates may be adjusted by between −4 to +6%, varying by taxon, although Kronenfeld [56] shows adjustments of less than 1%. These biases can be minimized by appropriate analytical decisions; many efforts over the years have assessed and corrected for the biases and idiosyncrasies in the original surveyor data [17,38,39,53,55,57–60]. And, even given these caveats, PLSS records remain the best source of data about both forest composition and structure in the United States prior to EuroAmerican settlement.

This analysis builds upon and merges previous state-level efforts to digitize and database the point-level PLSS data for Wisconsin, Minnesota and the Upper Peninsula and upper third of the Lower Peninsula of Michigan. These datasets were combined using spatial tools in R [61,62] to form a common dataset for the upper Midwest (Fig 1) using the Albers Great Lakes and St Lawrence projection (see code in Supplement 1, file: *step_one_clean_bind.R;* proj4: *+init:EPSG:3175*).

**Fig. 1.**
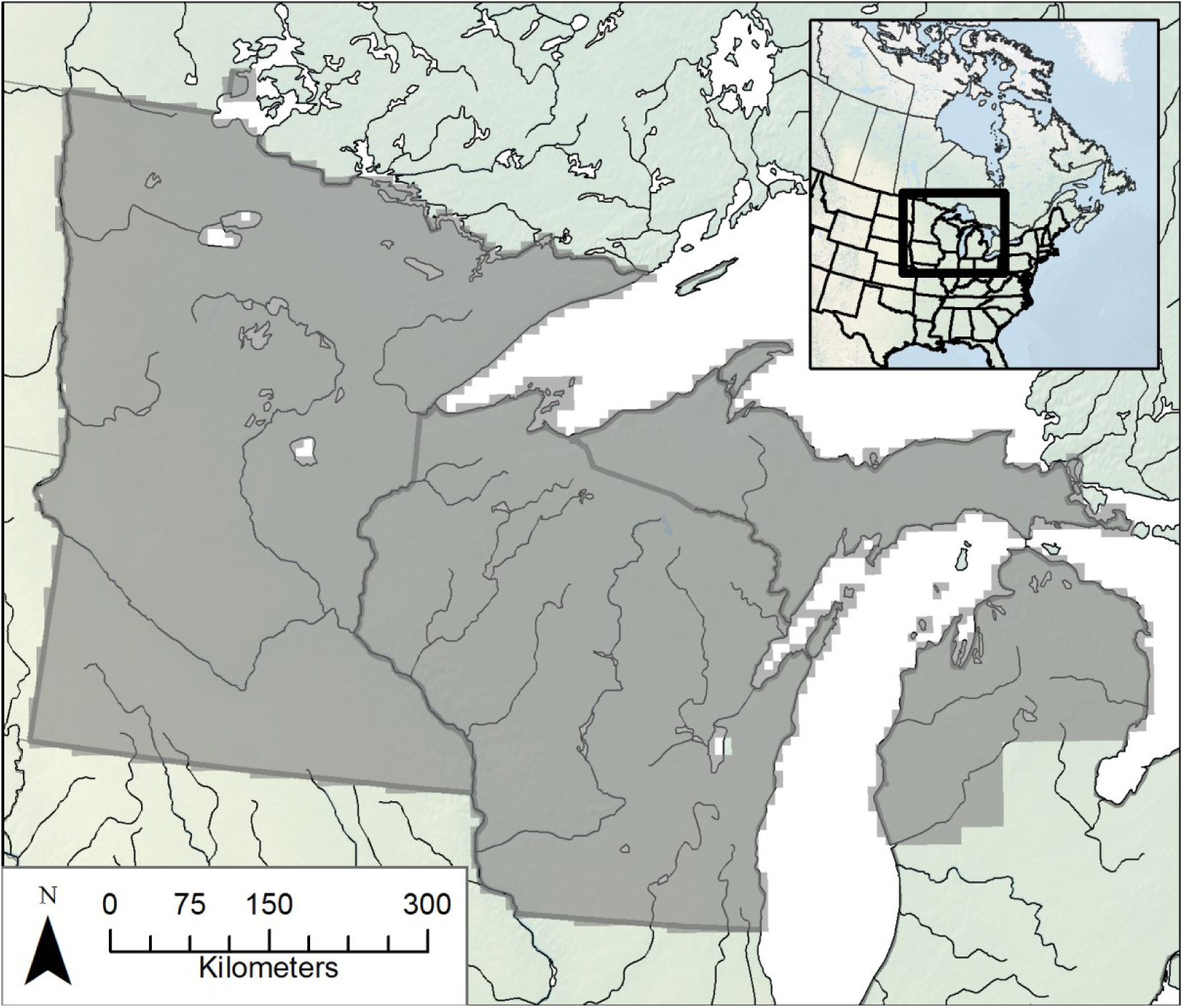
The domain of the Public Land Survey investigated in this study. The broad domain includes Minnesota, Wisconsin and the upper two thirds of Michigan state. A 8×8km grid is superimposed over the region to aggregate data, resulting in a total of 7940 cells containing data.

We took several steps to standardize the dataset and minimize the potential effects of surveyor bias upon estimates of forest composition, density, and biomass. All steps are preserved in the supplementary R code (Supplement 1: *step_one_clean_bind.R*). First, we excluded line and meander trees (i.e. trees encountered along survey lines, versus trees located at section or quarter corners) because surveyor selection biases appear to have been more strongly expressed for line trees, meander trees have non-random habitat preferences [38], and the inherent differences in sampling design between line, meander and corner points. We used only the closest two trees at each corner point because the third and fourth furthest trees have stronger biases with respect to species composition and diameter [38]. Corner points were used only if 1) there were at least two trees at a survey point, 2) the two trees were from different quadrants (defined by the cardinal directions), and 3) there were valid azimuths to the trees (a defined quadrant with an angle between 0 and 90) and valid diameters (numeric, non-zero).

Many species-level identifications used by PLSS surveyors are ambiguous. Statistical models can predict the identity of ambiguous species [51], but these models introduce a second layer of uncertainty into the compositional data, both from the initial surveyors’ identification, and from the statistical disambiguation. Given the regional scale of the analysis, and the inherent uncertainty in the survey data itself, we chose to avoid this layer of taxonomic uncertainty, and retained only genus-level identification (Supplement 2, *Standardized Taxonomy*). The ecological implications for the use of genera-level taxonomies are important for this region. While fire tolerance is fairly well conserved within genera, shade tolerance can vary. *Betula* contains shade intolerant *B. paperyfera* and the intermediate *B. alleghaniensis,* while *Pinus* contains the very shade intolerant *P. banksiana,* the intolerant *P. resinosa* and the shade tolerant *P. strobus.* For cases where shade tolerance (or fire tolerance) varies strongly within a genera we examine the data to determine the suitability of the assignment, or extent of confusion within the assigned genera.

In areas of open prairie or other treeless areas, *e.g.* southwestern Minnesota, surveyors recorded distances and bearings to ‘Non Tree’ objects. When points were to be located in water bodies the point data indicates ‘Water’. Points recorded “No Tree” are considered to have been from extremely open vegetation, with an estimated point-level stem density of 0 stems ha^-1^. We based our estimates on terrestrial coverage, so water cells are excluded completely. Hence, absence of trees at “No Tree” locations does reduce the gridded estimates of terrestrial stem density, but absence of trees at ‘Water’ locations does not.

Digitization of the original surveyor notebooks introduces the possibility of transcription errors. The Wisconsin dataset was compiled by the Mladenoff lab group, and has undergone several revisions over the last two decades in an effort to provide accurate data [30,38,43,44,51]. The Minnesota transcription error rate is likely between 1 and 5%, and the treatment of azimuths to trees varies across the dataset [37]. Michigan surveyor observations were transcribed to mylar sheets overlaid on State Quadrangle maps, so that the points were displayed geographically, and then digititized to a point based shapefile (Ed Schools, pers. comm.; Great Lakes Ecological Assessment. USDA Forest Service Northern Research Station. Rhinelander, WI. http://www.ncrs.fs.fed.us/gla/), carrying two potential sources of transcription error. Preliminary assessment of Southern Michigan data indicates a transcription error rate of 3 - 6%. To reduce errors associated with transcription across all datasets, we exclude sites for which multiple large trees have a distance of 1 link (20.12 cm) to plot center, trees with very large diameters (diameter at breast height - dbh > 100 in; 254 cm), plots where the azimuth to the tree is unclear, and plots where the tree is at plot center but has a recorded azimuth. All removed plots are documented in the code used for analysis (Supplement 1: *step_one_clean_bind.R)* and are commented for review.

### Data Aggregation

We binned the point data using an 64km^2^ grid (Albers Gt. Lakes St Lawrence projection; Supplement 1: *base_calculations.R*) to create a dataset that has sufficient numerical power for spatial statistical modeling and sufficient resolution for regional scale analysis [63]. This resolution is finer than the 100km^2^ gridded scale used in Freidman and Reich [45], but coarser than township grids used in other studies [19,56] to provide a scale comparable to aggregated FIA data at a broader scale. Forest compositional data is based on the number of individuals of each genus or plant functional type (PFT) present at all points within a cell. Stem density, basal area and biomass are averaged across all trees at all points within the cell.

### Stem Density

Estimating stem density from PLSS data is based on a plotless density estimator that uses the measured distances from each survey point to the nearest trees at the plot location [64,65]. The Morisita density estimator is then modified to minimize error due to different sampling geometries and several known surveyor biases [17,38,39,53,55,57–60]. The standardized approach for this method is well validated, however surveyors did not use a consistent approach to plot level sampling. Survey sampling instructions changed throughout the implementation of the PLSS in this region and differed between section and quarter section points and between internal and external points within a township [36,38]. Our approach allows for spatial variation in surveyor methods by applying various spatially different correction factors based not only on the empirical sample geometry, but also on known surveyor biases deviating from this design [57]. These estimates are based on empirical examination of the underlying data, and have been validated using simulations on stem mapped stands [57].

We estimate stem density (stems m^−2^) based on a on a modified form of the Morisita two-tree density estimator, which uses the distance-to-tree measurements for the two closest trees at each point [66]. Our modified form uses explicit and spatially varying correction factors, modeled after the Cottam correction factor [67], that account for variations in sampling designs over time and among surveyors. All code to perform the analysis is included in Supplement 1.

We estimate the basic stem density (stems m^−2^) using the point-to-tree distances for the closest trees to each point within a defined number of sectors around the point (Reference 64 eqn 31.):

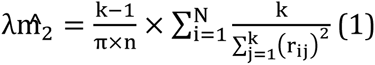

where λ is density ; k is the number of sectors within which trees are sampled, N is the number of points over which estimates are aggregated, r is the distance of point-to-tree (as m). This estimate can be modified by a refinement of the Cottam quadrant factors [66,67] which recognizes that different sampling designs, and the order of the distances in different quadrants (or sectors) carry specific weights. This correction, herein called κ, accounts for different sampling designs. When either four quadrants or trees are sampled (point quarter design), or when two trees in opposite semicircles (point halves design) are sampled, the equation is accurate and κ = 1; when the two trees are in the nearest of two quadrants (two nearest quadrants design), κ = 0.857; and when two trees are in quadrants on the same side of the direction of travel (one-sided or interior half design), κ = 2. This parameter, in Cottam’s notation [67], is a divisor of the denominator above, or here, the mathematically equivalent multiplier in the numerator of the reciprocal of the squared distances.

We further simplify the density estimate in equation (1) so that it is calculated at each point (N=1) and for two sample trees only (k=2):

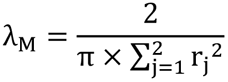

Then the point values for any sampling design can be Cottam corrected (κ × λ_M_). For example, the basic Morisita equation for two sectors assumes trees are located in opposite halves, so if the actual design is the nearest tree in the two nearest quadrants, the density from equation 2 will be overestimated and must be correspondingly corrected by multiplying by κ = 0.857.

Further corrections account for the restriction of trees to less than the full sector (θ), censoring of trees near the cardinal azimuths (ζ), and undersampling of trees smaller than a certain diameter limit (ϕ). These parameters are derived from analyses of measurements of bearing angles and diameters actually observed in surveys of witness trees within a subset of townships across the upper Midwest.

Sector bias (θ). Although the density model for two tree points assumes that the trees are on opposite sides of a sample line (point halves), the actual sample is often more restricted (< 180^°^) within the sector, or is a less restricted (> 180^°^) angle beyond the sector (see Supplement 3). This deviation from the equation’s assumption of equal distribution of angles across the 180^°^ sector is quantified using the empirical angle between the bearings of the two trees (pair angle). The pair angle frequencies (Supplement 3) that the observed proportion of trees (p) within any restricted sector divided by the proportion of that angle within the circle (α) are an estimate of the bias imposed by the actual sampling [55]. The factor (θ = p/α) indicates bias associated with differences in geometry of two tree samples. This parameter (θ) varies from 0.71 to 1.27, indicating sampling from effectively 253^°^ to 141^°^ sectors.

Azimuthal censoring (ζ). In addition to sector bias, surveyors did not always sample trees near the cardinal directions [55,58,59]. This azimuthal censoring is commonly found along the line of travel on section lines and sometimes on the perpendicular quarter-section lines. Trees near the cardinal directions were passed over, and a replacement was found within a more restricted angular region. The correction for this bias is calculated following Kronenfeld and Wang [55] in a manner similar to the sector bias. The factor ζ is the ratio of the proportion of trees in the restricted area (p) divided by the proportion of the complete circle (α) that is used. The azimuthal censoring parameter (ζ) ranges from 1.03 to 1.25 indicating an equivalent to complete elimination of trees from 10^°^ to 72^°^ azimuths adjacent to the cardinal directions.

Diameter limit (ϕ). Examination of the diameter distributions from settlement era surveys across the upper Midwest clearly demonstrate witness trees less than 8 inches in diameter were undersampled [38,57,59]. We have confirmed this bias in our own inspection of plots of diameter frequency in the PLSS data, which show a strong mode at 8”. This bias can be accommodated by setting a diameter limit, and only calculating the density for trees with diameters above this limit. Total density calculated from all trees is reduced to this reference limit by simply multiplying the total by the percentage of trees above this limit. This effectively eliminates the smaller trees from the total and normalizes the value of trees above this standard. The parameter (ϕ) represents diameter size bias is simply the percentage of trees ≥ 8” and, in practice, ranges from 0.6 - 0.9.

Because all surveyor bias corrections are simple multipliers of the model density and should be independent, the bias-minimized estimate of the point density of trees ≥ 8” is:

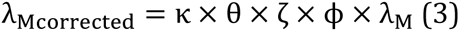

Estimates for each point *i* can be averaged for all *N* points in any region. Correction factors are calculated separately for different regions, years, internal versus external lines, section versus quarter-section points, and surveyor sampling designs (Supplement 4). All code to perform the analyses is included in Supplement 1 and the full rationale for and calculation of these measures is described further in Cogbill*el al.* [57]. Further, simulation used stem mapped stands from the region presented in Cogbill*el al.* [57] supports the robustness of this method, as opposed to other methods presented in the literature.

### Basal Area and Biomass Estimates

Forest basal area is calculated by multiplying the point-based stem density estimate by the average stem basal area from the reported diameters at breast height for the closest two trees at the point (n=2). Aboveground dry biomass (Mg ha^−1^) is calculated using the USFS FIA tree volume and dry aboveground biomass equations for the United States [68].

Biomass equations share the basic form:

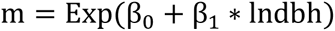

where m represents stem biomass for an individual tree in kg. β_0_ and β_1_ are parameters derived from [68] and described in Table 1. dbh is the stem diameter at breast height (converted to cm) recorded in the survey notes. The biomass estimates are summed across both trees at a survey point and multiplied by the stem density calculated at that point to produce an estimate of aboveground biomass reported in Mg ha^−1^ [68].

**Table 1.**
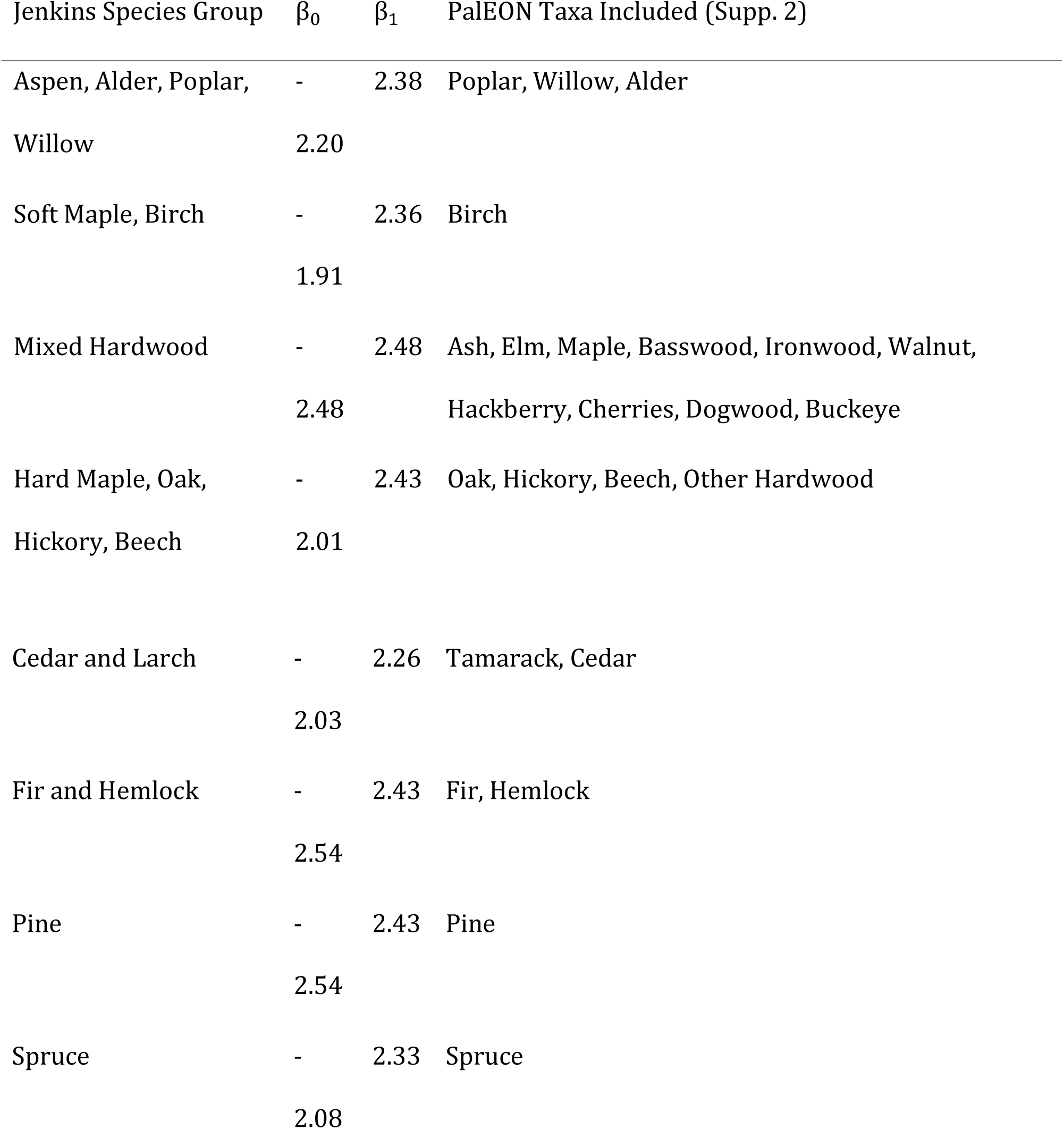
*Biomass parameters used for the calculation of biomass in the pre-settlement dataset(rounded for clarity).*

Matching PLSS tree genera to the species groups defined by Jenkins *et al.* [68] is straightforward, placing the 22 genera used in this study into 9 allometric groups (Table 1). However, all maples are assigned to the generic “Hardwood” group since separate allometric relationships exist for soft and hard maple (Table 1). Biomass estimates for “Non tree” survey points are assigned 0 Mg ha^−1^.

We use the stem density thresholds of Anderson and Anderson [69] to discriminate prairie, savanna, and forest.

### FIA Stem Density, Basal Area and Biomass

The United States Forest Service has monitored the nation’s forests through the FIA Program since 1929, with an annualized state inventory system implemented in 1998 [70]. On average there is one permanent FIA plot per 2,428 ha of land in the United States classified as forested. Each FIA plot consists of four 7.2m fixed-radius subplots in which measurements are made of all trees >12.7cm dbh [70]. We used data from the most recent full plot inventory (2007-2011). The FIA plot inventory provides a median of 3 FIA plots per cell using the 64km^2^ grid.

We calculated mean basal area (m^2^ ha^−1^), stem density (stems ha^−1^), mean diameter at breast height (cm) for all live trees with dbh greater than 20.32cm (8in). Biomass calculations (mean biomass: Mg ha^−1^) used the same set of allometric regression equations as for the PLSS data [68]. All calculations followed instructions in Woudenberg *et al* [70] using forested plots only (COND_STATUS_CD 1).

One critical issue is the reliance on forested condition for the FIA sampling. This reduces our capacity to compare forest state between PLS and FIA cover in regions with historical prairie and savanna coverage. In addition, it may result in the overestimation of modern density, basal area and biomass at the mesoscale in these same regions by drawing from a sample biased specifically towards regions with > 10% forest cover [70], however, the 10% cover theshold is fairly low, but more likely in line with “open forest” [69] than savanna.

### Gridding and Analysing PLSS and FIA Data

Maps of stem density, basal area and biomass were generated by averaging all PLSS point or FIA plot estimates within a 64km^2^ raster cell. Differences in sampling design between PLSS and FIA data combined with spatially structured forest heterogeneity will affect the partitioning of within-cell versus between-cell variance, but not the expected estimates. Most 64km^2^ cells have one or a few intensively sampled FIA plots. Therefore at this scale of aggregation, the low density of FIA plots in heterogeneous forests could result in high within-cell variance and high between-cell variability. For the PLSS plotless (point based) estimates, stem density estimates are sensitive to trees close to the plot center. Point-level estimates with very high stem densities can skew the rasterized values, and it is difficult to distinguish artifacts from locations truly characterized by high densities. To accommodate points with exceptionally high densities we carry all values through the analysis, but exclude the top 2.5 percentile when reporting means and standard deviations in our analysis. PLS-based estimates are highly variable from point to point due to the small sample size, but have low variance among 64 km^2^ raster cells due to the uniform sampling pattern of the data. Thus within-cell variance is expected to be high for the PLSS point data, but spatial patterns are expected to be robust at the cell level. The base raster and all rasterized data are available as Supplement 3.

Standard statistical analysis of the gridded data, including correlations, paired t-tests and regression, was carried out in R [61], and is documented in supplementary material that includes a subset of the raw data to allow reproducibility. Analysis and presentation uses elements from the following R packages: `cluster` [71], `ggplot2` [72,73], `gridExtra` [74], `igraph` [75], `mgcv` [76], `plyr` [77], `raster` [78], `reshape2` [79], `rgdal` [62], `rgeos` [80], `sp` [81,82], and `spdep` [83].

We identify analogs and examine differences in composition between and within PLSS and FIA datasets using Bray-Curtis dissimilarity [84] for proportional composition within raster cells using basal area measurements. For the analog analysis we are interested only in the minimum compositional distance between a focal cell and its nearest compositional (not spatial) neighbor. The distribution of compositional dissimilarities within datasets indicates forest heterogeneity within each time period, while the search for closest analogs between datasets indicates whether contemporary forests lack analogs in pre-settlement forests (‘novel forests’), or vice versa (‘lost forests’). For the analog analyses, we compute Bray-Curtis distance between each 64km^2^ cell in either the FIA or the PLSS periods to all other cells within the other dataset (FIA to FIA, PLSS to PLSS), and between datasets (PLSS to FIA and FIA to PLS), retaining only the minimum. For within era analyses (FIA - FIA and PLSS - PLSS), cells were not allowed to match to themselves. We define vegetation classes for lost and novel forests using k-medoid clustering [71].

The differences in sampling design and scale between the PLSS and FIA datasets, described above, potentially affect between-era assessments of compositional similarity [47]. The effects of differences in scale should be strongest in regions where there are few FIA plots per 64 km^2^ cell, or where within-cell heterogeneity is high. For the analog analyses, this effect should increase the compositional differences between the FIA and PLSS datasets. We test for the importance of this effect on our analog analyses via a sensitivity analysis in which we test whether dissimilarities between FIA and PLSS grid cells are affected by the number of PLSS plots per cell. We find a small effect (see below), suggesting that our analyses are mainly sensitive to the compositional and structural processes operating on large spatial scales.

To understand the extent to which the processes governing novelty operate at landscape scales, we relate the novelty of a cell to the spatial distance between individual novel cells and the nearest ‘remnant’ forest cell, *i.e.,* how far away can you go from a remnant forest cell before all cells are predicted to be novel. We examine whether this relationship varies between forest types, and whether it is different than the relationship we might see if the dissiminlarity values were distributed randomly on the landscape. The definition of “remnant” forest is likely to be arbitrary and, possibly, contentious. We use a threshold, the lowest 25%ile of compositional dissimilarity within the PLSS data, as our cutoff. This means that all FIA cells with nearest neighbor dissimilarities to the PLSS era forests below this cutoff are considered to be representative of the PLSS era forests. The analysis presented below is robust to higher cutoffs for the remnant forest threshold.

We use a generalized linear model with a binomial family to relate novelty (as a binomial, either novel or not) to the spatial distance from the nearest ‘remnant’ cell for each of the five major forest types within the PLSS data (Oak savanna, Oak/Poplar/Basswood/Maple, Pine, Hemlock/Cedar/Birch/Maple and Tamarack/Pine/Spruce/Poplar forests). Because the geographic extent of this region is complex, with islands, peninsulas and political boundaries, we use permutation, resampling the FIA to PLSS nearest neighbor distances without replacement, to estimate the expected distance to novelty if FIA to PLSS nearest neighbor dissimilarities were distributed randomly on the landscape.

We expect that a weak relationship will indicate that novelty, following landscape-scale land use change, is moderated by a species pool culled from small remnant patches, individual specimens, or local scale restoration efforts. A significant relationship between distance from remant forest and novelty indicates that small patches have been insufficient to restore natural forest cover within the region, and would indicate that greater efforts are needed to restore landscapes at regional scales.

All datasets and analytic codes presented here are publicly available and open source at (http://github.com/PalEON-Project/WitnessTrees), with the goal of enabling further analyses of ecological patterns across the region and the effects of post-settlement land use on forest composition and structure. Data are also archived at the Long Term Ecological Research Network Data Portal (https://portal.lternet.edu/nis/home.jsp).

## Results

### Data Standardization

The original PLSS dataset contains 490,818 corner points (excluding line and meander points), with 166,607 points from Wisconsin, 231,083 points from Minnesota and 93,095 points from Michigan Standardizing data and accounting for potential outliers, described above, removed approximately 1.5% points from the dataset, yielding a final total of 366,993 points with estimates used in our analysis.

Rasterizing the PLSS dataset to the Albers 64km^2^ grid produces 7,939 raster cells with data. Each cell contains between 1 and 94 corner points, with a mean of 61.8 (σ = 15) and a median of 67 corners (Supplement 3). Cells with a low number of points were mainly near water bodies or along political boundaries such as the Canadian/Minnesota border, or southern Minnesota and Wisconsin borders. Only 2.44% of cells have fewer than 10 points per cell.

Species assignments to genera were rarely problematic. Only 18 PLSS trees were assigned to the Unknown Tree category, representing less than 0.01% of all points. These unknown trees largely consisted of corner trees for which taxon could not be interpreted, but for which diameter and azimuth data was recorded. A further 0.011% of trees were assigned to the “Other hardwood” taxon (*e.g.,* hawthorn, “may cherry”, and “white thorn”).

For maple the class has very high within-genera specificity for a number of assignments. A total of 78478 trees are assigned to “Maple”. Of these, surveyors do use common names that can be ascribed to the species level (e.g., *A. saccharum,* n = 56331), but a large number of the remaining assignments are above the species level (n = 21356). This lack of specificity for a large number of records causes challenges in using the species level data. A similar pattern is found for pine, where many individual trees (125639) can be identified to the level of species (*P. strobus,* n = 41673; *P. banksiana,* n = 28784; *P. resinosa,* n = 28766), but there remains a large class of pine identified only at the genus level, or with unclear assignment (n = 17606).

For ash the data includes both surveyor attributions to “brown ash” (presumably a colloquial term used by surveyors as this is not currently an accepted common name in the region) and black ash (n=9312), and white ash (n = 2350), but again, also includes a large class of ash for which no distinction is made within the genera (n = 7423).

These patterns are repeated throughout the data. For spruce this within-genera confusion is even greater, with 50188 assignments to genera-level classes and only 20 to either black or white spruce.

### Spatial Patterns of Settlement-Era Forest Composition: Taxa and PFTs

#### Stem Density, Basal Area and Biomass

The mean stem density for the region (Fig 2a) is 153 stems ha^−1^. Stem density exclusive of prairie is 172 stems ha^−1^ and is 216 stems ha^−1^ when both prairie and savanna are excluded. The 95th percentile range is 0 - 423 stems ha^−1^, and within-cell standard deviations between 0 and 441 stems ha^−1^. Basal area in the domain (Fig 2c) has a 95th percentile range between 0 and 63.5 m^2^ ha^−1^, a mean of 22.2 m^2^ ha^−1^, within-cell standard deviations range from 0 to 76.7 m^2^ ha^−1^. Biomass ranges from 0 to 209 Mg ha^−1^ (Fig 2d), with cell level standard deviations between 0 and 569 Mg ha^−1^. High within-cell standard deviations relative to mean values within cells for density, basal area and biomass indicate high levels of heterogeneity within cells, as expected for the PLSS data, given its dispersed sampling design.

**Fig. 2.**
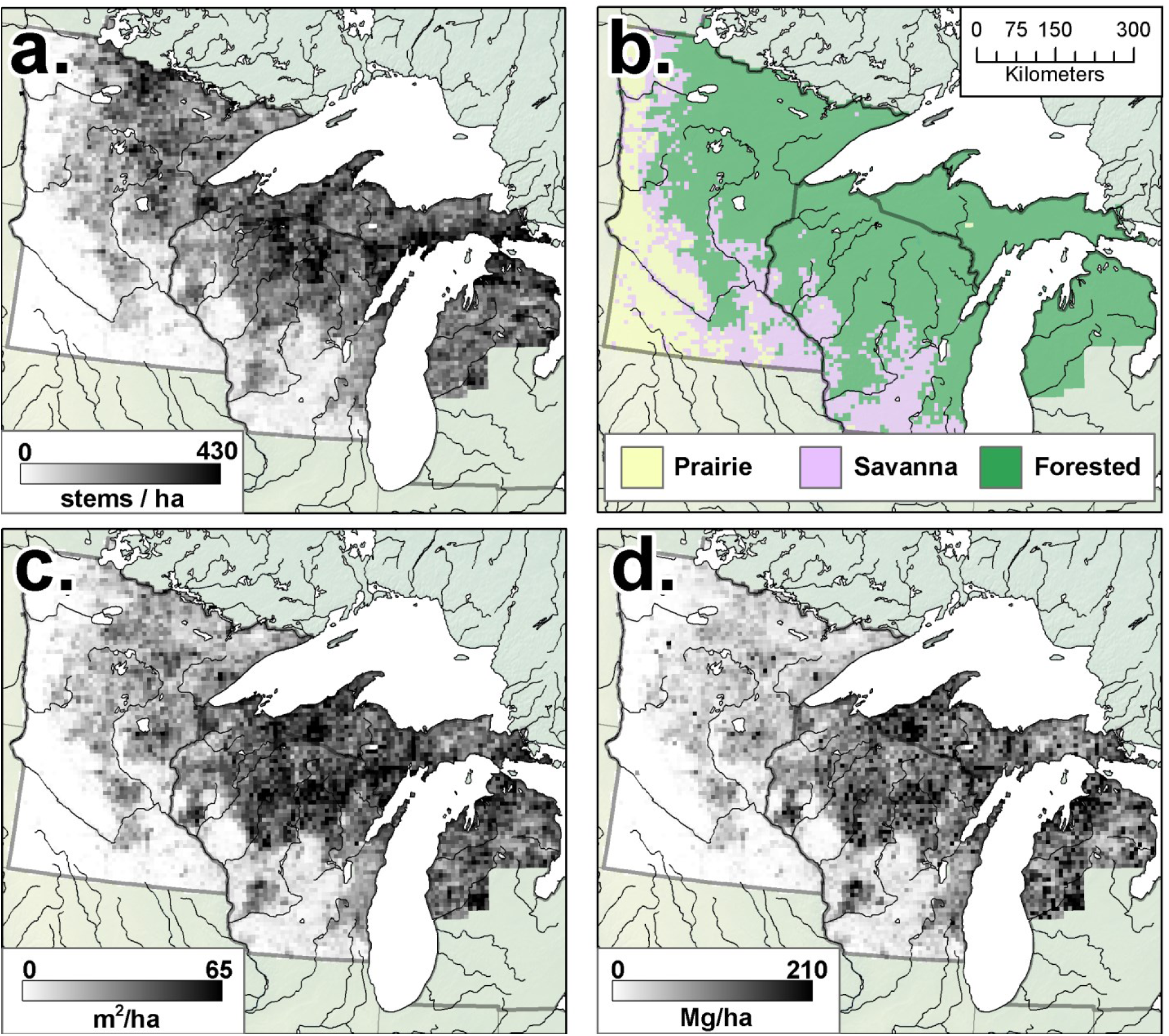
*Total stem density (a) in the Upper Midwest, along with forest type classification (b) based on PLSS data and the stem density thresholds defined by Anderson and Anderson [69]; Table 2). Fine lines represent major rivers. To a first order, basal area (c) and biomass (d) show similar patterns to stem density (but see* Fig 3).

In the PLSS data, stem density is lowest in the western and southwestern portions of the region, regions defined as prairie and savanna (Fig 2b, Table 2). When the Anderson and Anderson [69] stem density thresholds (<47 stems ha^−1^ for Savanna, Table 2) are used, the extent of area classified as savanna is roughly equivalent to prior reconstructions [22,85,86] (Fig 2b). The highest stem densities occur in north-central Minnesota and in north-eastern Wisconsin (Fig 2a), indicating younger forests and/or regions of lower forest productivity.

**Table 2.** *Forest classification scheme used in this paper for comparison between presettlement forests and the Haxeltine and Prentice [87] potential vegetation classes represented in Ramankutty and Foley [1]. Plant functional types (PFTs) for grasslands (CG, grassland; Non-Tree samples in the PLS), broad leafed deciduous taxa (BDT) and needleleaded evergreen taxa (NET) are used, but leaf area index used in Haxeltine and Prentice [87] is replaced by stem density classes from Anderson and Anderson [69].*

| Forest Class | Haxeltine & Prentice Rules | Current Study |
| --- | --- | --- |
| Prairie | Dominant PFT CG, LAI > 0.4 | Stem dens. < 0.5 stem/ha |
| Savanna | Dominant PFT CG, LAI > 0.6 | $1 < \text{Stem dens.} < 47 \text{ stems } \text{ha}^{-1}$ |
| Temperate | Dominant PFT BDT, LAI > 2.5 | Stem dens. > 48 stems $\text{ha}^{-1}$, BDT > 70% |
| Deciduous |  | composition |
| Temperate | Dominant PFT (NET + NDT), | Stem dens. > 47 stems ha<sup>-1</sup>, NET + NDT |
| Conifer | LAI > 2.5 | > 70% composition |
| Mixedwood | Both BDT (LAI > 1.5) & NET<br>(LAI > 2.5) present | Stem dens. > 47 stems ha<sup>-1</sup>, BDT & NET<br>both < 70% composition |

Forest structure during the settlement era can be understood in part by examining the ratio of stem density to biomass, a measure that incorporates both tree size and stocking. Regions in northern Minnesota and northwestern Wisconsin have low biomass and high stem densities (Fig 3, blue). This indicates the presence of young, small-diameter, even-aged stands, possibly due to frequent stand-replacing fire disturbance in the pre-EuroAmerican period or to poor edaphic conditions. Fire-originated vegetation is supported by co-location with fire-prone landscapes in Wisconsin [88]. High-density, low-biomass regions also have shallower soils, colder climate, and resulting lower productivity. High-biomass values relative to stem density (Fig 3, red) are found in Michigan and southern Wisconsin. These regions have higher proportions of deciduous species, with higher tree diameters than in northern Minnesota.

**Fig. 3.**
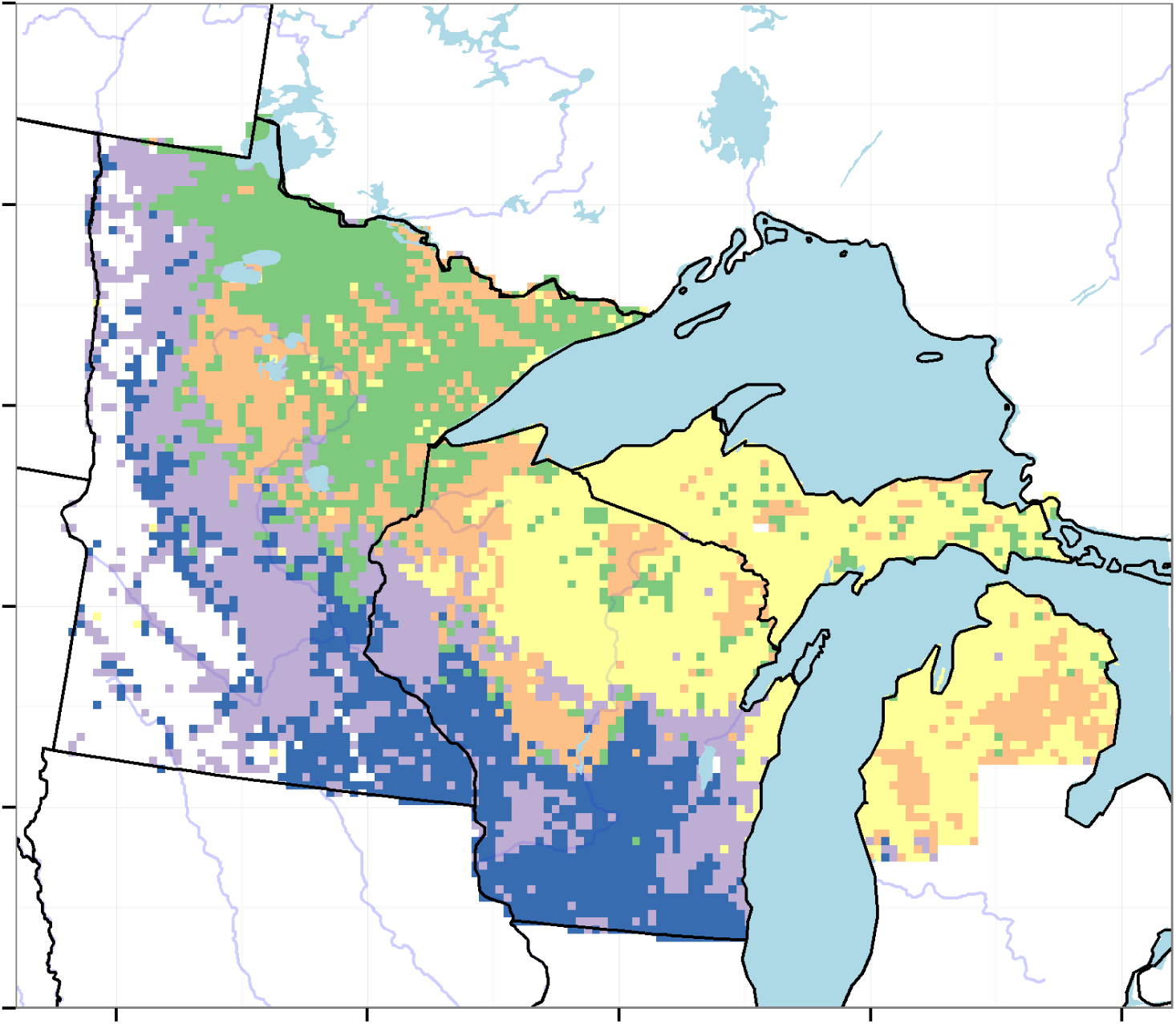
*The major forest types in the pre-settlement Upper Midwest. Five clusters are shown using k-medoid clustering. These clusters represent (b) the ratio between biomass and stem density as an indicator of forest structure. Regions with high stem density to biomass ratios (blue) indicate dense stands of smaller trees, while regions with low stem density to biomass ratios (red) indicate larger trees with wider spacings.*

Taxon composition within settlement-era forests is spatially structured along dominant gradients from south to north (deciduous dominated to conifer dominated forests) and from east to west (mixed wood forests to open prairie) (Fig 4). Oak is dominant in the south of the region, with an average composition of 21%, however, that proportion drops to 8% when only forested cells (cells with stem density > 48 stems/ha) are considered, due to its prevalence as a monotypic dominant in the savanna and prairie. Pine shows the opposite trend, with average composition of 14% and 17% in unforested and forested cells respectively. Pine distributions represent three dominant taxa, *Pinus strobus, Pinus resinosa* and *Pinus banksiana.* These three species have overlapping but ecologically dissimilar distributions, occuring in close proximity in some regions, such as central Wisconsin, and are typically associated with sandy soils with low water availability. Other taxa with high average composition in forested cells include maple (10%), birch (10%), tamarack (9%) and hemlock (8%).

**Fig. 4.**
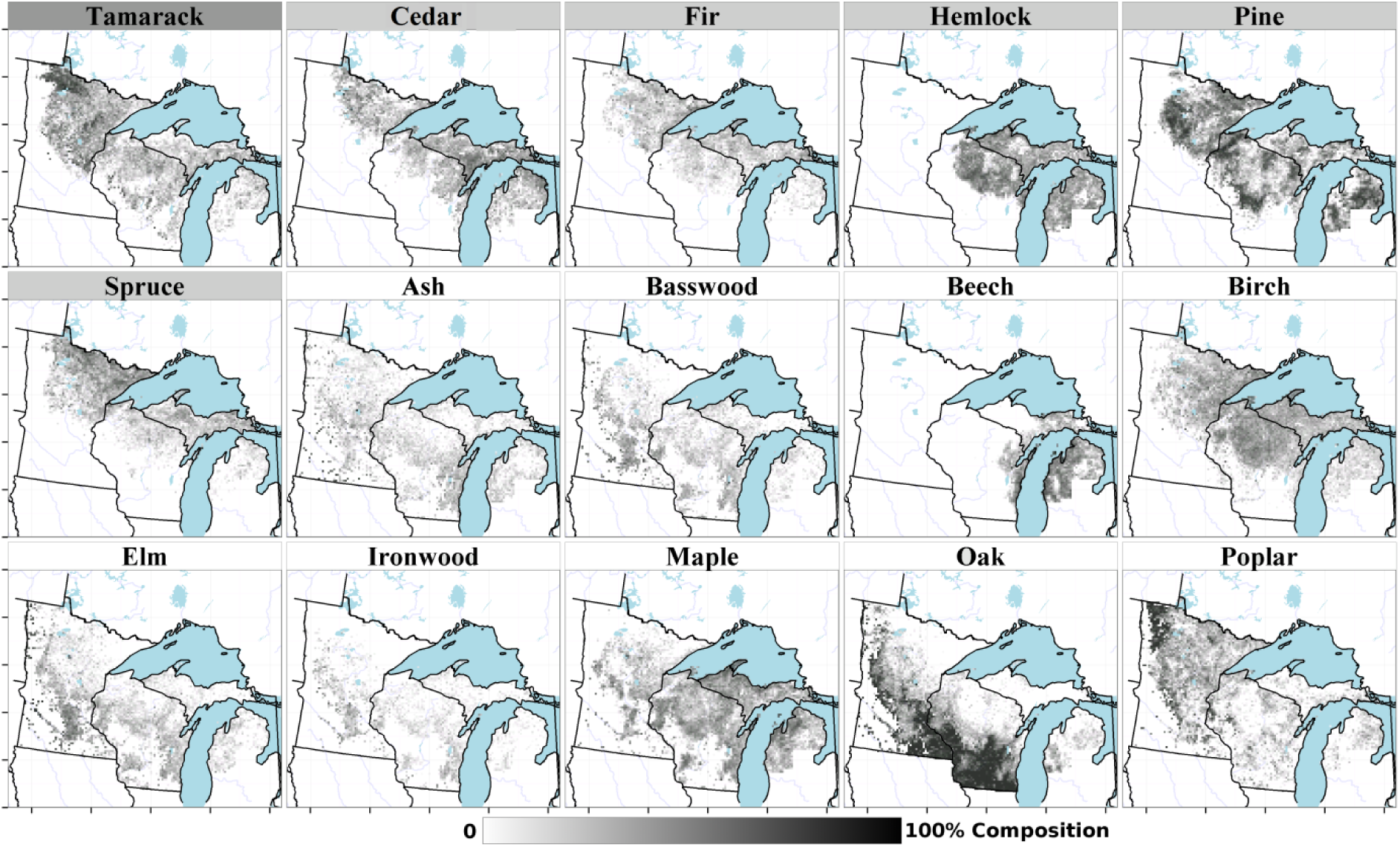
*Forest composition (%) for the 15 most abundant tree taxa. The scale is drawn using a square-root transform to emphasize low abundances. Shading of the bar above individual taxon maps indicates plant functional type assignments (dark gray: needleleafed deciduous; light gray: needleleafed evergreen; white: broadleafed deciduous).*

For a number of taxa, proportions are linked to the total basal area within the cell. For 4 taxa - hemlock, birch, maple and cedar - taxon proportions are positively related to total basal area. For 17 taxa including oak, ironwood, poplar, tamarack and elm, high proportions are strongly associated with lower basal areas (Figures 3 and 5). This suggests that hemlock, birch, maple and cedar occurred in well-stocked forests, with higher average dbh. These taxa are most common in Michigan and in upper Wisconsin. Taxa with negative relationships to total basal area (*e.g*., spruce and tamarack) are more common in the northwestern part of the domain.

Spruce in the PLSS represents two species (*Picea glauca, Picea mariana*) with overlapping distributions, but complex site preferences that vary in space. *P. glauca* is generally associated with dry upland to wet-mesic sites, while *P. mariana* is associated with hydric sites, but *P. mariana* also frequently occupies upland sites in northern Minnesota. Both cedar (*Thuja occidentalis*) and fir (*Abies balsamea*) are mono-specific genera in this region.

Northern hardwoods, such as yellow birch and sugar maple, and beech, are much less common in the lower peninsula of Michigan, and southern Wisconsin, except along Lake Michigan. Birch has extensive cover in the north, likely reflecting high pre-settlement proportions of yellow birch (*Betula alleghaniensis*) on mesic soils, and paper birch on sandy fire-prone soils and in northern Minnesota (birch proportions reach upwards of 34.1% in northeastern Minnesota). Hardwoods in the southwest, such as oak, elm, ironwood and basswood, are most typically mono-specific groupings, with the exception of oak, which comprises 7 species (see Supplement 2). Hardwoods in the southwest are located primarily along the savanna and southern forest margins, or in the southern temperate deciduous forests. Finally, maple and poplar (aspen) have a broad regional distribution, occupying nearly the entire wooded domain. Poplar comprises four species in the region, while maple comprises five species (Supplement 2). Both hardwood classes, those limited to the southern portions of the region, and those with distributions across the domain, correspond to well-defined vegetation patterns for the region [85].

These individual species distributions result in a mosaic of forest classes accross the region (Fig 5). The dominant class is the Hemlock/Cedar/Birch/Maple assemblage in northern Wisconsin, and upper Michigan (Fig 5, yellow). This mixedwood assemblage is interspersed by both Pine dominated landscapes (Fig 5, orange) and, to a lesser degree, the softwood assemblage Tamarack/Pine/Spruce/Poplar (Fig 5, green), which dominates in northeastern Minnesota. The softwood assemblage is itself interspersed with Pine dominated landscapes, and grades into a mixed-hardwood assemblage of Oak/Poplar/Basswood/Maple (Fig 5, light purple) to the west. Thismixed-softwood forest assemblage grades south into mono-specific Oak savanna (Fig 5, dark blue).

**Fig. 5.**
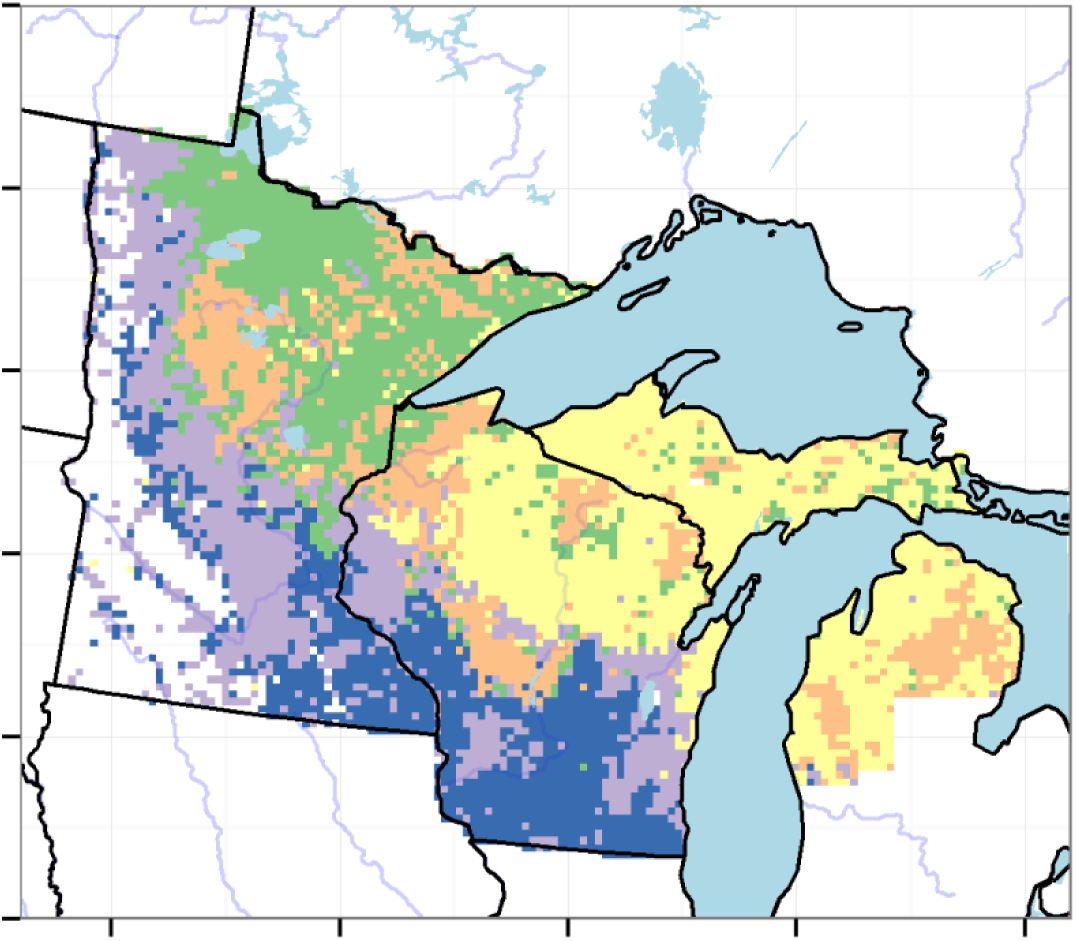
*The five dominant forest types in the Upper Midwest as defined by k-medoid clustering. Forest types (from largest to smallest) include Hemlock/Cedar/Birch/Maple (yellow), Oak/Poplar/Basswood/Maple (light purple), Tamarack/Pine/Spruce/Poplar (light green), Oak Savanna (dark purple) and Pine (orange). These forest types represent meso-scale (64km^2^) forest associations, rather than local-scale associations.*

The broad distributions of most plant functional types results in patterns within individual PFTs that are dissimilar to the forest cover classes (Fig 5). Thus overlap among PFT distributions (Fig 6) emerges from the changing composition within the plant functional type from deciduous broadleaved species associated with the southern, deciduous dominated region, to broadleafed deciduous species associated with more northern regions in the upper Midwest.

**Fig. 6.**
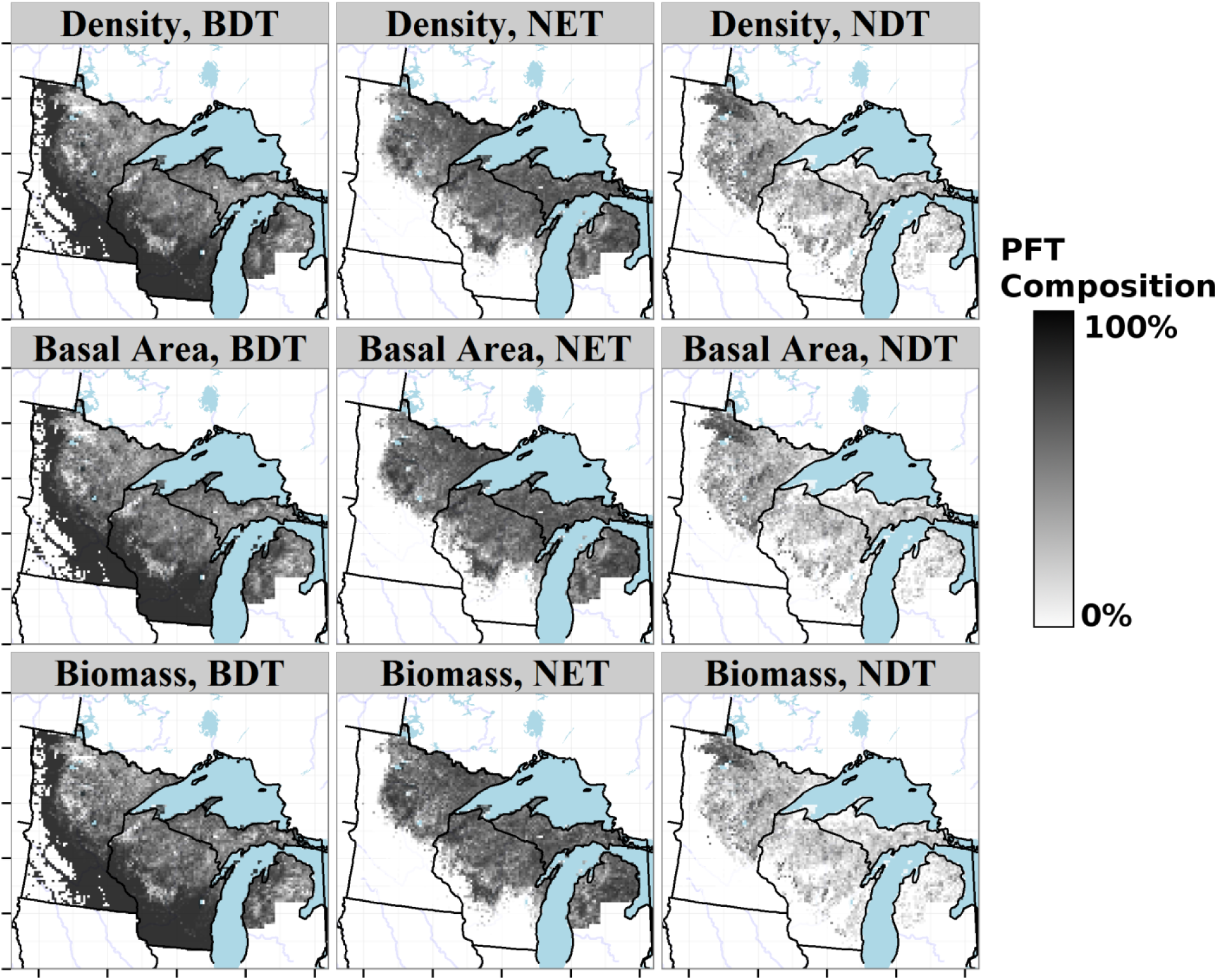
*Proportional distribution of Plant Functional Types (PFTs) in the upper Midwest from PLSS data, for broadleaved deciduous trees (BDT), needleleaved deciduous trees (NDT), and needleleaved evergreen trees (NET). Distributions are shown as proportions relative to total basal area, total biomass, and composition* (Fig 2*). The grassland PFT is mapped onto non-tree cells with the assumption that if trees were available surveyors would have sampled them.*

### Structural Changes Between PLSS and FIA Forests

Differences between PLSS and FIA data shows strong spatial patterns, but overall estimates can be examined. By cell, modern forests (FIA) have higher stem densities (146 stems ha^−1^, t_1,5177_ = 51.8, p < 0.01) than PLSS forests, but slightly lower basal areas (−4.5 m^2^ ha^−1^, t_1,5177_ = −16.4, p < 0.01) and lower biomass (−8.72 Mg ha^−1^, t_1,5177_ = −6.55, p < 0.01) (Fig 7). We use only point pairs where both FIA and PLSS data occur since non-forested regions are excluded from the FIA and as such cannot be directly compared with PLS estimates. The similarity in biomass despite lower stem density and total basal area in the PLSS data is surprising. Two likely factors are shifts in allometric scaling associated with changes in species composition, or a higher mean diameter of PLSS trees (Fig 7d). Total biomass was 45,080 Mg higher in the PLSS when summed across all cells coincident between the FIA and PLSS.

**Fig. 7.**
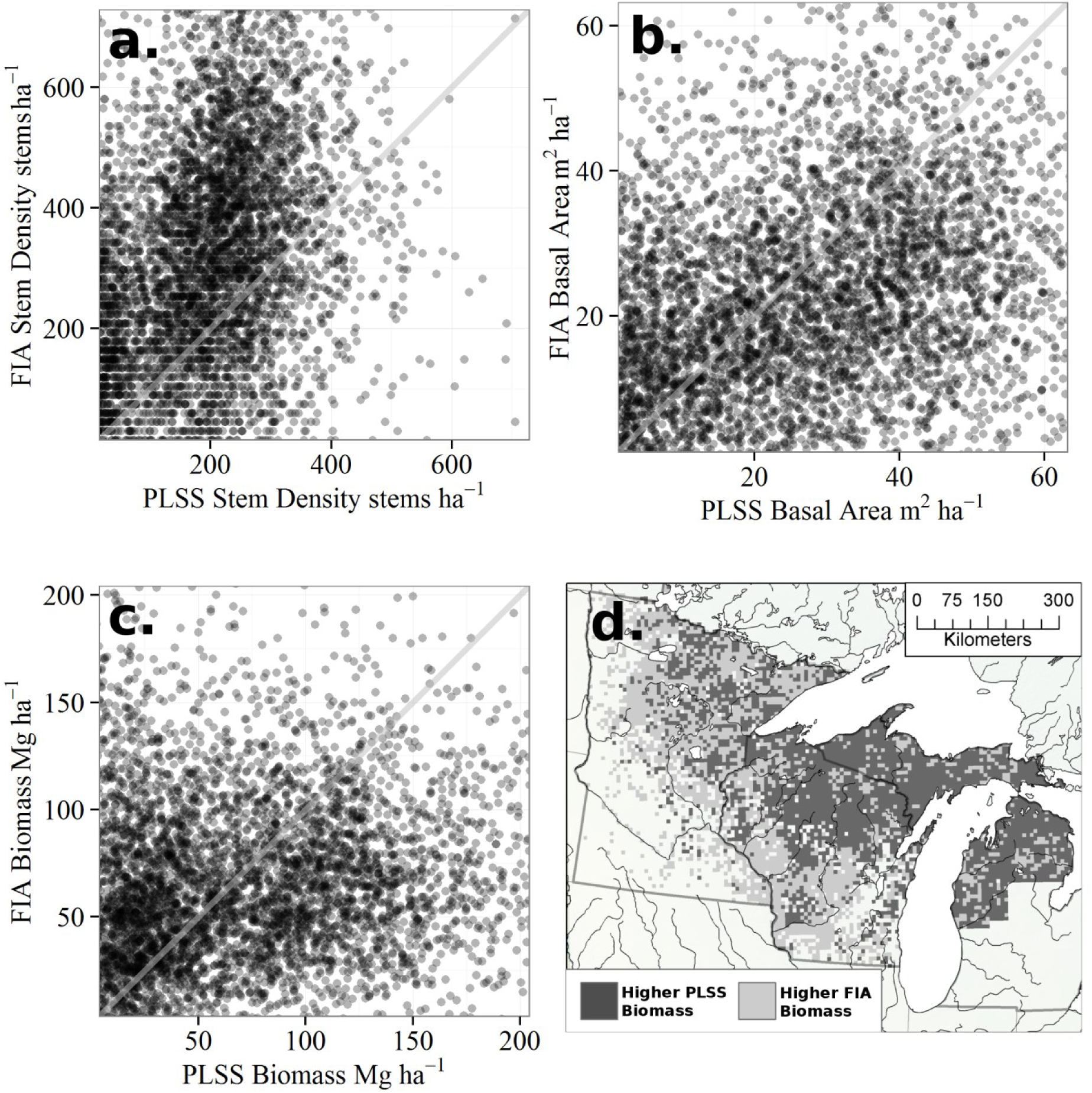
*The relationship between (a) average stem density, (b) total basal area and (c) biomass values in the PLSS and FIA datasets. Stem density and total basal area are higher in the FIA than in the PLS, however average cell biomass is higher in the PLSS. A 1:1 line has been added to panels a-c to indicate equality.*

Every one of the five historical PLSS zones shows an increase in stem density (Table 3). The two forest types bordering the prairie, Oak Savanna and Oak/Poplar/Basswood/Maple both show increases in density that likely reflect, in part, the issues addressed earlier with regards to the sampling of forested plots in the FIA (over 10% cover). Density in the Oak Savanna increases from a mean 27 stems/ha to 217 stems/ha, with a mean biomass increase of 62 Mg ha^−1^ per cell (Table 3), the highest of any of the zones. The Oak/Poplar/Basswood/Maple forest had higher PLSS-era densities (90 stems/ha) reflecting open forest status rather than savanna (Table 2), but also shows a large increase in estimated FIA-era stem density (to 218 stems/ha) but with a much lower increase in biomass than the Oak Savanna, and a negligable increase in basal area (Table 3). The largest forest zone, Hemlock/Cedar/Birch/Maple shows the largest decline in biomass (a net loss of 56.7 MG ha^01^ since the PLSS-era) and basal area (net loss of 13 m^2^ ha^−1^ since the PLSS-era), but with an average increase in FIA era stem density.

**Table 3.**
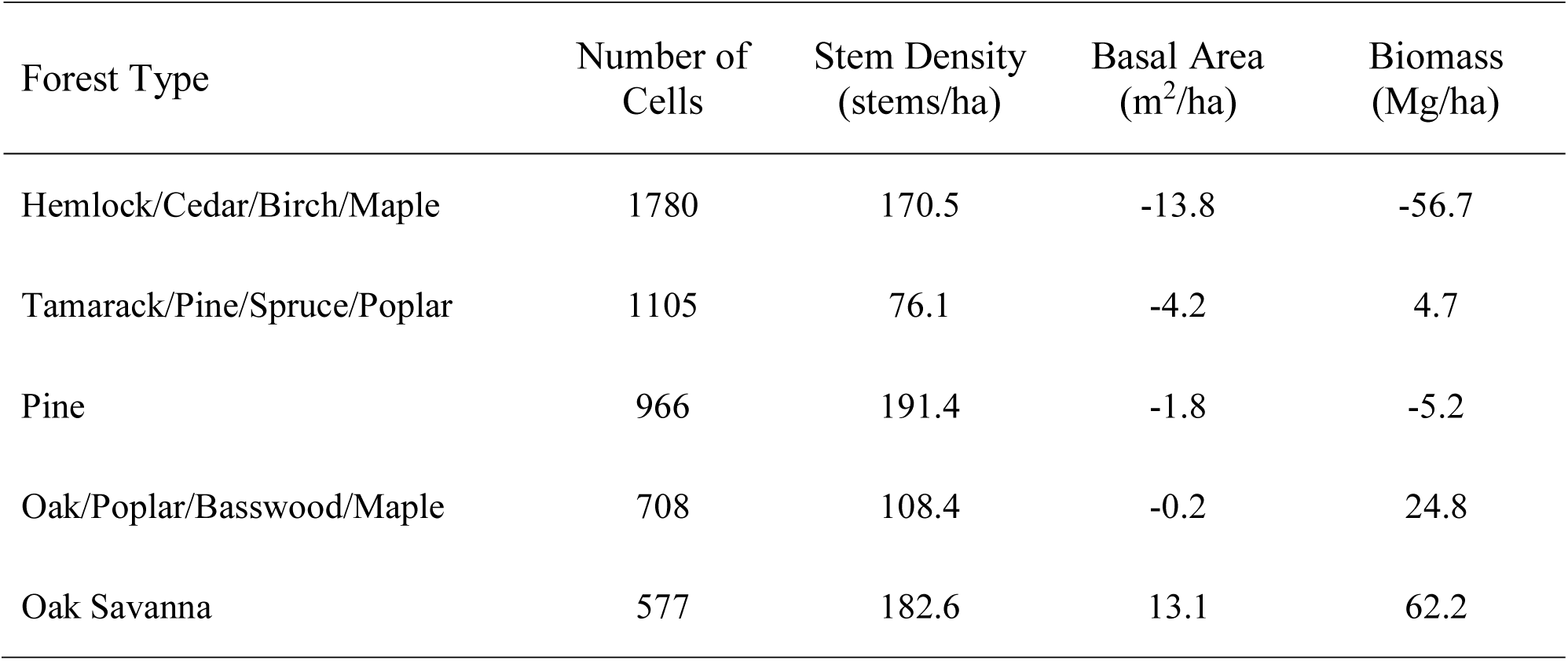
*Mean cell-wise change in forest zone density, basal area and biomass since PLSS for cells with coverage in both PLSS and FIA eras by forest class. All forest zones show increases in stem density since the PLSS era (positive values). All zones but Oak Savanna most show declines in mean basal area since the PLSS era, while modern biomass is lower in the FIA-era for the both Hemlock/Cedar/Birch/Maple and Pine forest zones, but higher in the remaining three zones.*

| Forest Type | Number of Cells | Stem Density (stems/ha) | Basal Area (m<sup>2</sup>/ha) | Biomass (Mg/ha) |
| --- | --- | --- | --- | --- |
| Hemlock/Cedar/Birch/Maple | 1780 | 170.5 | -13.8 | -56.7 |
| Tamarack/Pine/Spruce/Poplar | 1105 | 76.1 | -4.2 | 4.7 |
| Pine | 966 | 191.4 | -1.8 | -5.2 |
| Oak/Poplar/Basswood/Maple | 708 | 108.4 | -0.2 | 24.8 |
| Oak Savanna | 577 | 182.6 | 13.1 | 62.2 |

The PLSS has a lower overall mean diameter than the FIA (*δ*_diam_ = −2.9 cm, 95%CI from - 17.3 to 8.18cm). FIA diameters are higher than PLSS diameters in the northwestern parts of the domain (on average 6.47 cm higher), overlapping almost exactly with regions where we have shown low biomass-high density stands (Fig 3). At the same time, regions with high biomass and low density stands, in northeastern Wisconsin, and the Upper and Lower Peninsulas of Michigan, had higher average diameters during the PLSS than in the FIA, on average 3.65 cm higher. Thus we are seeing an overal increase in tree size in the sub-boreal region and a decrease in temperate mixedwood forests, where we find tree species with much higher dbh:biomass ratios [68]. This is coupled with declining variance in dbh across the domain (from within cell variance of 37.9 cm the PLSS to 30.7 cm in the FIA). Thus, the mechanism by which low density and basal area produce roughly equivalent biomass estimates between the FIA and PLSS is likely due to the differential impacts of land clearence and subesequent forest management in the south east vs the northwest. The loss of high biomass southern hardwood forests is balanced by higher biomass in the northeast due to fire suppression and regeneration of hardwoods in the northwest. Declining diameters from the PLSS to FIA are most strongly associated with higher abundances of poplar, ironwood and oak, while declining diameters are associated with maple and hemlock, further supporting the assertion that much of the loss in diameter, and, subsequently biomass, is occuring in southeastern mixedwood/hardwood forest, while diameter and biomass increases are occuring in the northwest.

Differences between FIA and PLSS data in sampling design are unlikely to be a factor for most measures (see below); these differences are expected to affect how these datasets sample local- to landscape-scale heterogeneity, but should not affect the overall trends between datasets. Differences in variability introduce noise into the relationship, but given the large number of samples used here, the trends should be robust.

### Compositional Changes Between PLSS and FIA Forests: Novel and Lost Forests

Both the PLS- and FIA-era compositional data show similar patterns of within-dataset dissimilarity, with the highest dissimilarities found in central Minnesota and northwestern Wisconsin. High within-PLSS dissimilarities are associated with high proportions of maple, birch and fir while high within-FIA dissimilarities are associated with high proportions of hemlock, cedar and fir. Dissimilarity values in the FIA dataset are less spatially structured than in the PLSS. Moran’s I for dissimilarities within the FIA (I_FIA_ = 0.198, p < 0.001) are lower than the dissimilarities within the PLSS (I_PLSS_ = 0.496, p < 0.001), suggesting lower spatial autocorrelation in the FIA dataset. Cells with identical pairs represent 5.6% of the PLSS cells and 7.44% of FIA cells. Identical cells in the PLSS are largely located along the southern margin and most (69.5%) are composed entirely of oak. Cells in the FIA with identical neighbors are composed of either pure oak (19.4%), pure poplar (26%) or pure ash (14%).

There is a small but significant positive relationship (F_1,5964_ = 9 2 0, p < 0.001) between the number of FIA plots and within-FIA dissimilarity. The relationship accounts for 13% of total variance and estimates an increase of δ_d_ = 0.0134 for every FIA plot within a cell. This increase represents only 3.08% of the total range of dissimilarity values for the FIA data. There is a gradient of species richness that is co-linear with the number of FIA plots within a cell, where plot number increases from open forest in the south-west to closed canopy, mixed forest in the Upper Peninsula of Michigan. Hence, differences in within- and between-cell variability between the PLSS and FIA datasets seem to have only a minor effect on these regional-scale dissimilarity analyses.

We define no-analog communities as those whose nearest neighbour is beyond the 95%ile for dissimilarities within a particular dataset. In the PLSS dataset, forests that have no modern analogs are defined as “lost forests”, while forest types in the FIA with no past analogs are defined as “novel forests”. More than 25% of PLSS sites have no analog in the FIA dataset (‘lost forests’; PLS-FIA dissimilarity, Fig 8c), while 29% of FIA sites have no analog in the PLSS data (‘novel forests’; FIA-PLSS dissimilarity, Fig 8d). Lost forests show strong spatial coherence, centered on the “Tension Zone” [85], the ecotone between deciduous forests and hemlock-dominated mixed forest (Fig 4).

**Fig. 8.**
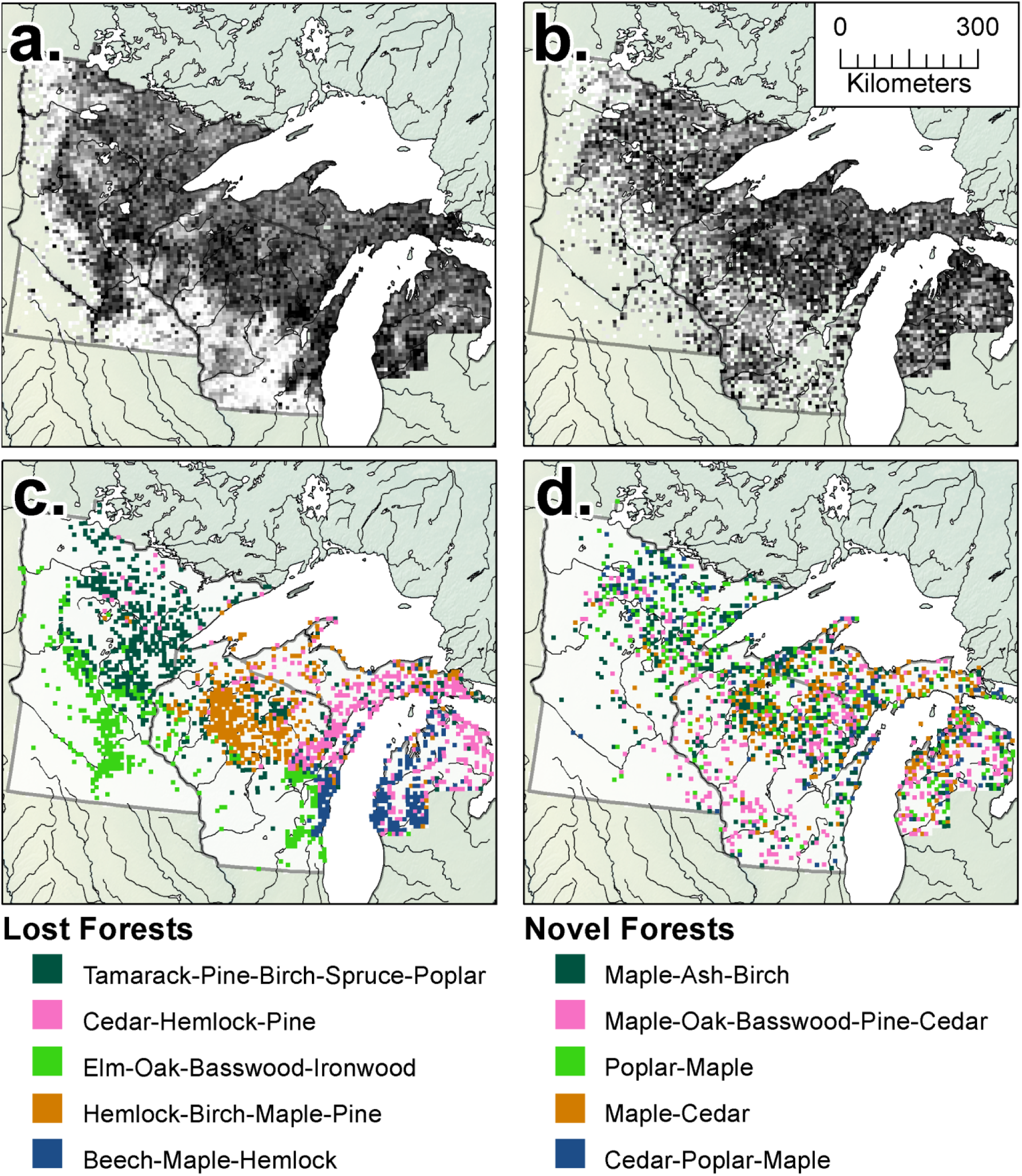
*Minimum dissimilarity maps. Distributions of minimum (within dataset) dissimilarities during the PLSS (a) and FIA (b) show spatially structured patterns of dissimilarity, with stronger spatial coherence for the PLS. Lost forests (c) show strong compositional and spatial coherence, and have more taxa with percent composition > 10% than within Novel forests during the FIA era (d).*

Lost forests are drawn from across the domain, and show strong ecological and spatial coherence (Fig 8c). Forest classes generally fall into five classes: Tamarack-Pine-Birch-Spruce-Poplar accounts for 28.8% of all lost forests and 7.97% of the total region. This forest type is largely found in north eastern Minnesota, extending southward to central Minnesota, into Wisconsin and along the Upper Peninsula of Michigan, as well as in scattered locations on the Lower Peninsula of Michigan (Fig 8c). This forest likely represents a mesic to hydric forest assemblage, particularly further eastward. Modern forests spatially overlapping this lost type are largely composed of poplar 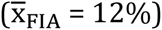 and oak 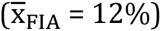. Tamarack in these forests has declined significantly, from 23% to only 5% in the FIA, while Poplar has increased from 10% to 22%, resulting in forests that look less mesic and more like early seral forests.

Cedar/juniper-Hemlock-Pine accounts for 19.8% of all lost forests and 5.49% of the total region. This forest type is found largely in northeastern Wisconsin, and the Upper and Lower Peninsulas of Michigan. This lost forest type has been predominantly replaced by maple, poplar, and pine, retaining relatively high levels of cedar 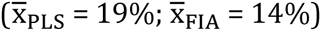. The loss of hemlock is widespread across the region, but particularly within this forest type, declining to only 3% from a pre-settlement average of 18%.

Elm-Oak-Basswood-Ironwood accounts for 19.6% of all lost forests and 5.42% of the total region. The region is centered largely within savanna and prairie-forest margins, both in south-central Minnesota and in eastern Wisconsin, but, is largely absent from savanna in the Driftless area of southwestern Wisconsin. These forests were historically elm dominated 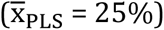, not oak dominated savanna, as elsewhere (particularly in the Driftless). Modern forests replacing these stands are dominated by oak and ash, with strong components of maple, and basswood. Elm has declined strongly in modern forests 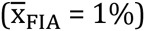, possibly in part due to Dutch Elm Disease and land use. The increase in ash in these forests is substantial, from 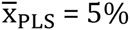 to 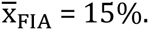

Hemlock-Birch-Maple-Pine accounts for 19.2% of all lost forests and 5.33% of the total region. This forest type, dominant in north central Wisconsin, was dominated by hemlock 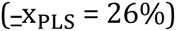 and what was likely late seral yellow birch 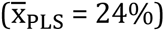, replaced largely by maple (from 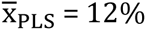 to 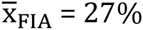). Poplar increases from 1% to 13% in the FIA, again indicating a shift to earlier seral forests in the FIA. Hemlock is almost entirely lost from the forests, declining from 26% to 4% in the FIA.

Lastly, Beech-Maple-Hemlock accounts for 12.6% of all lost forests and 3.49% of the total region. This forest type is found exclusively on the central, western shore of Lake Michigan and in the Lower Peninsula, in part due to the limited geographic range of Beech in the PLSS dataset (Fig 4). Beech is almost entirely excluded from the modern forests in this region, declining from 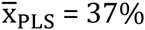 to 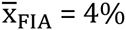. Pine in the region increases from 9% to 16%, while maple, the dominant taxa in the modern forests, increases from 16 - 25%.

On average lost forests contain higher proportions of ironwood (r = 0.203), beech (r = 0.2), birch (r = 0.189) and hemlock (r = 0.188) than the average PLSS forest, and lower proportions of oak (r = -0.28), poplar (r = -0.145), and pine (r = -0.107).

The distribution of novel ecosystems (Fig 8d) is spatially diffuse relative to the lost forest of the PLSS and the forest types tend to have fewer co-dominant taxa. FIA novel forest types also have a more uneven distribution in proportion than the PLSS lost forests. Overall, novel forests are associated with higher proportions of maple (r = 0.02), ash (r = 0.03) and basswood (r = -0.04), although basswood is dominant in only one forest type (Poplar-Cedar/juniper-Maple). Novel forests are associated with lower proportions of oak (r = −0.28), and pine (r = −0.11). This analysis suggests that the loss of particular forest types associated with post-settlement land use was concentrated in mesic deciduous forests and the ecotonal transition between southern and northern hardwood forests, while the gains in novelty were more dispersed, resulting from an overall decline in seral age.

By far the largest novel forest type is Maple, which accounts for 37.2% of all novel forests and 2.68% of the total region. As with all novel forest types, this forest type is broadly distributed across the region. This forest type is associated with co-dominant maple 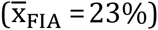 and ash 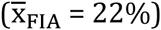. Hemlock has declined significantly across this forest type, from 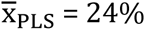 to 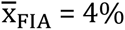.

Poplar-Cedar/juniper-Maple, accounts for 28.8% of all novel forests and 2.08% of the total region. The broad distributiof these novel forests makes assigning a past forest type more difficult than for the PLSS lost forests, the distribution replaces two classes of past forest, one dominated by oak, in southern Wisconsin and Minnesota, the other by mixed hemlock, beech, birch and cedar forests.

Pine-Cedar/juniper-Poplar-Maple forest accounts for 16.3% of all novel forests and 1.17% of the total region. This forest type is again broadly distributed, and is widely distributed across the region, representing a homogenous, early seral forest type, likely associated with more mesic sites. Oak forest accounts for 13.3% of all novel forests and 0.96% of the total region. This grouping again shows a pattern of broad distribution across the region, associated with cedar/juniper percentages near 40%, with smaller components of poplar (14%) and maple (13%).

### Spatial Correlates of Novelty

Modern compositional dissimilarity from the PLSS data is related to distance from ‘remnant’ forest. The dissimilarity quantile of FIA-PLSS distances increases with increasing distance to remnant cells, and this relationship is robust to higher thresholds for remnant forest classification, up to the 90%ile of within-PLSS near neighbor dissimilarities. Using the 25%ile for within PLSS dissimilarity, approximately 67% of FIA cells can be classed as ‘remnant’ forest. The mean distance to remnant forests for cells with dissimilarities above the 25%ile is 17.7 km, higher than the mean of ~9.6km expected if each 8x8km cell had at least one adjacent ‘remnant’ cell.

The GLM shows that distance from remnant forests in the FIA is significantly related to the probability of a cell being novel (χ_1_,_4_=623, p < 0.001). The mean distance to novelty varies by PLSS forest type, but is between approximately 20 and 60km for the four forest types examined here (Fig 9), while the null model would predict a distance of 10 - 20km to novelty from remnant cells if dissimilarities were distributed randomly on the landscape (Table 4). Novel forests are generally further from remnant patches than expected in the null model, regardless of forest type, but the distance to novelty is greater for modern forests that are, generally, more similar to their PLSS state (Pine and Tamarack dominated forests), and closer for forests that are more dissimilar.

**Fig. 9.**
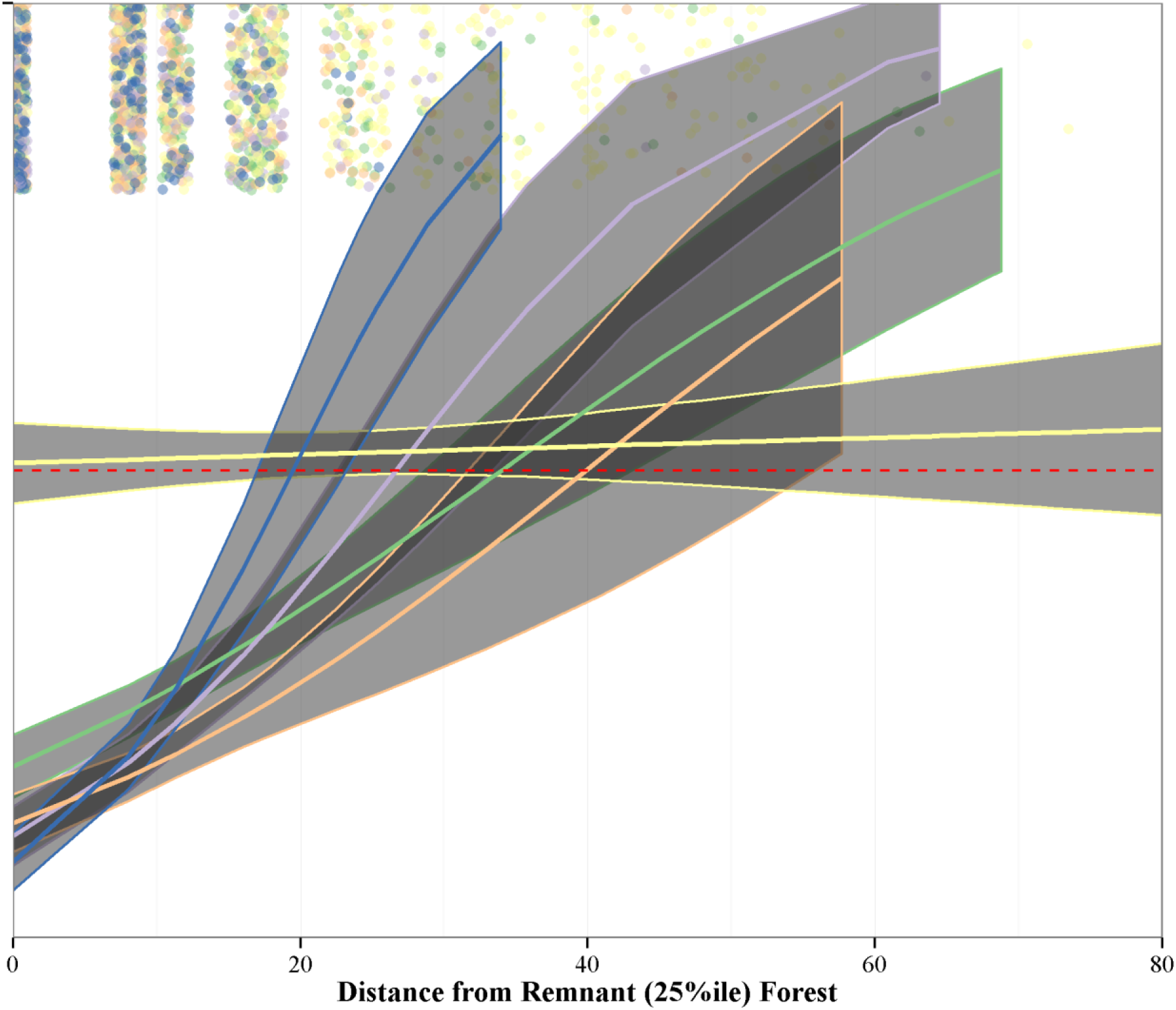
*The model relating novelty to spatial distance from remnant forest. Here the 25%ile is used to indicate remnant forest, and the 95%ile is defined as novelty. We use a binomial regression to predict novelty, the red dashed line indicates a response greater than 0.5. The curves represent the relationship between spatial distance and compositional dissimilarity for each of the five major historic forest types (Fig 5) defined here as Oak Savanna (blue), Oak/Poplar/Basswood/Maple (light purple), Tamarack/Pine/Spruce/Poplar (green), Hemlock/Cedar/Birch/Maple (yellow) and Pine (orange).*

**Table 4.** *Spatial distance to novelty - modeled as a binomial - from remnant forests (forests within the first 25th percentile of nearest neighbor distances). The null model uses permutation (n=100) where quantiles are resampled without replacement.*

| Zone | Min | Max | Min (Null) | Max (Null) |
| --- | --- | --- | --- | --- |
| Tamarack/Pine/Spruce/Poplar | 29 | 43 | 11 | 14 |
| Oak/Poplar/Basswood/Maple | 23 | 33 | 14 | 20 |
| Pine | 32 | 56 | 10 | 12 |
| Hemlock/Cedar/Birch/Maple | 0 | undef. | 11 | 13 |
| Oak Savanna | 17 | 23 | 13 | 18 |

Critically, we see that the Hemlock/Cedar/Birch/Maple forest class (Fig 5 & 10b, yellow), appearing as a flat line, predicts novelty continuously, from distance 0. This is due, in part, to the very small proporion of Hemlock/Cedar/Birch/Maple cells that are considered residual (only 63 of 1780 cells in the Hemlock zone are considered remnant) and the very high proportion of novel cells in the zone (923 of 1780 cells, or 52% of all cells).

Oak Savanna is the most similar to its null model, with a confidence interval that overlaps slightly with the null expectation (Table 4). Northern softwood forests (Tamarack/Pine/Spruce/Poplar, Fig 5, light green) reach novelty at between 29 and 43km, northern Oak forests (Oak/Poplar/Basswood/Maple; Fig 5, light purple) reach novelty at 23 - 33 km, slightly higher than the 14 - 19km predicted by the null model. Pine forests (Fig 5, orange) are three times further than expected by the null, at 32 - 56km (Table 4).

### Compositional Changes Between PLSS and FIA Forests: Ecotone Structure

To understand how the ecotonal structure has been transformed by post-settlement land use, we constructed two transects of the FIA and PLSS data (Fig 10a), and fitted GAM models to genus abundances along these transects. Transect One (T1) runs from northern prairie (in northern Minnesota) to southern deciduous savanna in southeastern Wisconsin (left to right in Figures 11c-f), while Transect Two (T2) runs from southern prairie in southwestern Minnesota to northern mixedwood forest in the Upper Peninsula of Michigan (left to right in Figures 11g-j). In general, these transect analyses show: 1) significant differences in ecotonal structure between the present and pre-settlement, and 2) steeper ecotones in the past and more diffuse ecotones today.

**Fig. 10.**
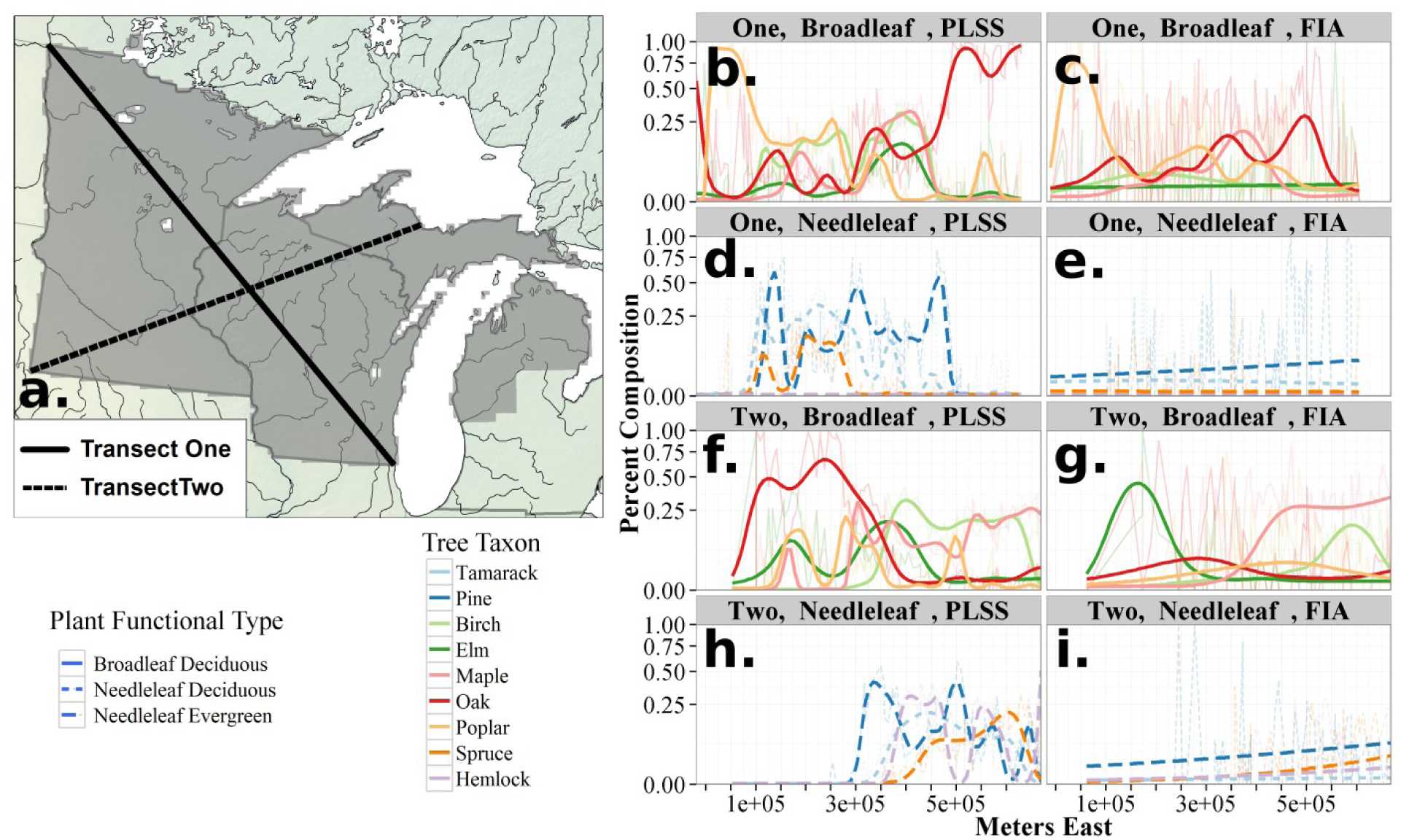
*Transects (a) across the region show clear changes in the ecotonal strength. Transect One shows shifts in broad-leafed taxon distributions from the PLSS to FIA (b and c) and in needle-leafed distributions (d and e). Transect Two broadleaf (f and g) and needleleaf (h and i) taxa show shifts that again appear to represent regional scale homogenization. Ecotones in the pre-settlement era were stronger in the past than they are in the present. Fitted curves represent smoothed estimates across the transects using Generalized Additive Models using a beta family.*

For T1, GAM models show significant differences (using AIC) between time periods in curves for all broadleafed taxa (Fig 10b & c) and for all needleleafed taxa (Figures 10d and e). The PLSS curves show a rapid transition in the northwest from oak to poplar dominated open forest that then transitions to a needleleafed forest composed of pine, spruce and tamarack, with high proportions of tamarack grading to pine further to the south east. Tamarack and poplar proportions decline gradually from the east, being replaced first by pine, then briefly by maple and birch, and then. ultimately by oak as the transect grades into oak savanna. In the FIA dataset oak and poplar in the northwest appears to decline simultaneously, grading into needleleafed forests that are absent from the FIA dataset in the first 100km along the transect. While the PLSS transect shows distinct vegetation types in the central partof the transect, the FIA shows relatively constant proportions of oak, pine, spruce, poplar and maple before pine, oak and elm increase in the southeastern portions of the transect.

The second transect shows a similar pattern, with well defined ecotones in the pre-settlement period(Fig 10f and h), that are largely absent from the FIA data (Fig 10g and i). Oak forest, with a component of elm and poplar in the southwest grades slowly to a rapid transition zone where pine, elm, maple (first), then rapidly birch, hemlock and tamarack, and later, spruce, increase. This region, the Tension Zone, extends from 3 × 10^5^ to 4.5×10^5^ meters East, eventually becoming a forest that shows co-dominance between birch, pine, maple, spruce and tamarack, likely reflecting some local variability as a result of topographic and hydrological factors. Missing data at the beginning of the FIA transect reflects a lack of FIA plots in unforested regions in the west

Contemporary forests show broader homogenization and increased heterogeneity (evidenced by the lower within-FIA Moran’s I estimates for near-neighbor distances) at a local scale in the region. Homogenization is evident across T1, where Bray-Curtis dissimilarity between adjacent cells declines from the PLSS to the FIA (δ_beta_ = −0.22, t_113_ = − 7.93, p<0.001), mirroring declines in the pine barrens between the 1950s and the present [18]. The PLSS shows strong differentiation in the central region of T2 where maple-pine-oak shifts to pine-poplar-birch forest (Fig 10d). This sharp ecotone is not apparent in the FIA data, which shows gradual and blurred changes in species composition across the ecotone (Fig 10i). β-diversity along T2 is lower in the FIA than in the PLSS (δ_beta_ = −0.19, t_65_=−7.34, p < 0.01), indicating higher heterogeneity in the PLSS data at the 64 km^2^ mesoscale.

Across the entire domain, β diversity is lower in the FIA than in the PLSS (δ_β_ = −0.172, t_1.3e7_ = 2480, p <0.001), lending support to the hypothesis of overall homogenization. Differences in sampling design between PLSS and FIA data cannot explain this homogenzation, since its effect would have been expected to increase β-diversity along linear transects and at larger spatial scales.

## Discussion

Many forests of the PLS, are no longer a part of the modern landscape. Forest types have been lost at the 64 km^2^ mesoscale, and new forest types have been gained. The joint controls of broad-scale climatic structuring and local hydrology on forest composition and density can be seen in the pre-settlement forests, particularly along the Minnesota River in south-western Minnesota, where a corridor of savanna was sustained in a region mostly occupied by prairie (Fig 2b), but ecotones in the modern forest composition data are weaker now than in the past (Fig 10), with clear signs of increased homogenization at local and regional scales and decreased spatial structure in vegetation assemblages (Fig 8).

The loss of ecotones in the upper Midwestern United States suggests that our ability to predict the abiotic controls on species distributions at the landscape scale may be weaker than in the past, reducing the influence of variables such as climate or edaphic factors, and increasing the relative influence of recent land use history. Our results suggest that both recent land use history and historical vegetation cover play a large role in recovery from the large scale disturbance seen following EuroAmerican settlement.

Work in eastern North America suggests the utility of including spatial structure in species distribution models to improve predictive ability [89]. The spatial random effects may improve models by capturing missing covariates within SDMs [89], but if recent land use history has strongly shaped species distributions, or co-occurence, then the spatial effect is likely to be non-stationary at longer temporal scales. Given the implicit assumption of stationarity in many ecological models [21], the need for longer time-scale observations, or multiple baselines from which to build our distributional models becomes critical if we are to avoid conflating recent land use effects with the long term ecological processes structuring the landscape.

Decreased β diversity along regional transects indicates homogenization at meso-scales of 100s of km^2^, while the overall reduction in Moran’s I for dissimilarity in the FIA indicates a regional reduction in heterogeneity on the scale of 1000s of km^2^. The selective loss or weakening of major vegetation ecotones, particularly in central Wisconsin, and the development of novel species assemblages across the region further suggests that modern correlational studies, examining regional relationships between species and climate (for example) may fail to capture the full range of edaphic controls on spcies distributions.

These changes are the result of land use, both agricultural and logging, but affect forests in contrasting ways across the domain. Maple has become one of the most dominant taxa across the region, while in northern Minnesota, forest biomass has increased and species shifts have reflected increases in poplar and pine, while in central and eastern Wisconsin, biomass has declined, and hemlock has been lost almost completely.

Anthropogenic shifts in forest composition over decades and centuries seen here and elsewhere [2,48] are embedded within a set of interacting systems that operate on multiple scales of space and time [90]. Combining regional historical baselines, long term ecological studies and high frequency analyses can reveal complex responses to climate change at local and regional scales [91]. Estimates of pre-settlement forest composition and structure are critical to understanding the processes that govern forest dynamics because they represent a snapshot of the landscape prior to major EuroAmerican land-use conversion [38,52]. Pre-settlement vegetation provides an opportunity to test forest-climate relationships prior to land-use conversion and to test dynamic vegetation models in a data assimilation framework [92]. For these reason, the widespread loss of regional forest associations common in the PLSS (Fig 8d), and the rapid rise of novel forest assemblages (Fig 8e) have important implications for our ability to understand ecological responses to changing climate. The loss of historical forest types implies that the modern understanding of forest cover, climate relationships, realized and potential niches and species associations may be strongly biased in this region, even though 29% of the total regional cover is novel relative to forests only two centuries ago.

Beyond shifts in composition at a meso-scale, the broader shifts in ecotones can strongly impact models of species responses and co-occurence on the landscape. For example, the heterogeneity, distribution, and control of savanna-forest boundaries [93] is of particular interest to ecologists and modelers given the ecological implications of current woody encroachment on savanna ecosystems [94]. Declines in landscape heterogeneity may also strongly affect ecosystem models, and predictions of future change. Our data show higher levels of vegetation heterogeneity at mesoscales during the pre-settlement era, and greater fine scaled turnover along transects. Lower β diversity shown here and elsewhere [18] indicate increasing homogeneity at a very large spatial scale, and the loss of resolution along major historical ecotones.

This study also points to the need for a deeper understanding of some of the landscape- and regional-scale drivers of novelty, given the likely role for climatic and land use change (including land abandonment) to continue to drive ecological novelty [95,96]. In particular the role of regional species pools and remnant patches of forest in driving or mitigating compositional novelty. This work shows that the baseline forest type, and its structure on the landscape moderates the degree to which landscape scale patterns can drive compositional novelty. To some degree relationships between compositional novelty and distance from remnant patches may be dependent on the simplicity or complexity of the species pool and the sensitivity of dissimilarity metrics to β diversity [97]. Our results indicate that diversity alone cannot be the driving factor in determining post-settlement dissimilarity (and novelty), since all forest classes show this pattern of change.

Both Pine and the Oak/Poplar/Basswood/Maple forest types are the most fragmented across the region. There is strong evidence that in some locations pine forests have persisted over very long timescales in the region [98], although there is also evidence, in other regions, that these states may shift strongly in response to interactions between landscape level processes such as fire and geophysical features [99]. Thus complex interactions between landscape scale processes, whether they be fire, land use change, or geophysical features, and the species assemblages themselves, point to the difficulty in making simplifying assumptions about species assemblages. Caution in simplifying species assignments, whether they be plant functional types, species richness, or phylogenetic metrics, is neccessary since this region is dominated by forests that respond very differently to the settlement-era (and pre-settlement) disturbance, but that are composed of different species of the same genera and plant functional type. This caution is clearly warranted since recent ecosystem model benchmarking using pre-settlement vegetation has shown significant mismatch between climate representations of plant functional types across a range of ecosystem models, with no model accurately representing the true climate space of plant functional types in the northeastern upper Midwestern United States [100].

The analysis relating to the distance-to-novelty (Fig 9) points to the possibility that landscape-scale restoriation has high likelihood of success if local-scale restoration focuses on sites where restoration potential is high, as suggested for Hemlock/Cedar/Birch/Maple forests in northern Wisconsin [86]. If some of the novelty is driven by depauparate species pools beyond certain threshold distances from remnant forests then it should also be possible to restore these forest at a regional scale through the translocation of key species [101]. This work is supported by a number of other studies at smaller scales [102–104], for example, the presence of white pine in mesic sites during the PLS era has been attributed to its presence as a seed source on marginal sites at scales of of hundreds of meters [105]. Computer simulations [106] show that seed source distribution can affect community composition over hundreds of years at large spatial scales in a region spatially coincident with this current study. Thus land use change has significantly altered the landscape, both by “resetting” the sucessional clock, but also, because of the extent of change, by impacting the regional species pool and seed source for re-establishing forests that are compositionally similar to pre-settlement forests.

Methodological advances of the current work include 1) the systematic standardization of PLSS data to enable mapping at broad spatial extent and high spatial resolution, 2) the use of spatially varying correction factors to accommodate variations among surveyors in sampling design, and 3) parallel analysis of FIA datasets to enable comparisons of forest composition and structure between contemporary and historical time periods. This approach is currently being extended to TPS and PLSS datasets across the north-central and northeastern US, with the goal of providing consistent reconstructions of forest composition and structure for northeastern US forests at the time of EuroAmerican forests.

Our results support the consensus that robust estimates of pre-settlement forest composition and structure can be obtained from PLSS data [39,44,46,107,108]. Patterns of density, basal area and biomass are roughly equivalent to previous estimates [16,19], but show variability across the region, largely structured by historical vegetation type (Table 3). Our results for stem density are lower than those estimated by Hanberrry *et al.* [17] for eastern Minnesota, but density and basal area are similar to those in the northern Lower Peninsula of Michigan [109] and biomass estimates are in line with estimates of aboveground carbon for Wisconsin [19].

These maps of settlement-era forest composition and structure can provide a useful calibration dataset for pollen-based vegetation reconstructions for time periods prior to the historic record [110]. Many papers have used calibration datasets comprised of modern pollen samples to build transfer functions for inferring past climates and vegetation from fossil pollen records [111–114]. However, modern pollen datasets are potentially confounded by recent land use, which can alter paleoclimatic reconstructions using pollen data [113]. By linking pollen and vegetation at modern and historical periods we develop capacity to provide compositional datasets at broader spatio-temporal scales, providing more data for model validation and improvement. Ultimately, it should be possible to assimilate these empirical reconstructions of past vegetation with dynamic vegetation models in order to infer forest composition and biomass during past climate changes. Data assimilation, however, requires assessment of observational and model uncertainty in the data sources used for data assimilation. Spatiotemporal models of uncertainty are being developed for the compositional data [63].

Ultimately the pre-settlement vegetation data present an opportunity to develop and refine statistical and mechanistic models of terrestrial vegetation that can take multiple structural and compositional forest attributes into account. The future development of uncertainty estimates for the data remains an opportunity that can help integrate pre-settlement estimates of composition and structure into a data assimilation framework to build more complete and more accurate reconstructions of past vegetation dynamics, and to help improve predictions of future vegetation under global change scenarios.

## Acknowledgements

The authors would like to thanks the large number of individuals who have worked to first, undertake the PLS survey, to bring the original survey data together, to digitize and standardize much of the survey results, and finally, to assist in interpreting and compiling the data in its present form. We would like to thank our reviewers and those who have sent comments on the preprint (http://dx.doi.org/10.1101/026575).

